# Population-level variation of enhancer expression identifies novel disease mechanisms in the human brain

**DOI:** 10.1101/2021.05.14.443421

**Authors:** Pengfei Dong, Gabriel E. Hoffman, Pasha Apontes, Jaroslav Bendl, Samir Rahman, Michael B. Fernando, Biao Zeng, James M. Vicari, Wen Zhang, Kiran Girdhar, Kayla G. Townsley, Ruth Misir, the CommonMind Consortium, Kristen J. Brennand, Vahram Haroutunian, Georgios Voloudakis, John F. Fullard, Panos Roussos

## Abstract

Identification of risk variants for neuropsychiatric diseases within enhancers underscores the importance of understanding the population-level variation of enhancers in the human brain. Besides regulating tissue- and cell-type-specific transcription of target genes, enhancers themselves can be transcribed. We expanded the catalog of known human brain transcribed enhancers by an order of magnitude by generating and jointly analyzing large-scale cell-type-specific transcriptome and regulome data. Examination of the transcriptome in 1,382 brain samples in two independent cohorts identified robust expression of transcribed enhancers. We explored gene-enhancer coordination and found that enhancer-linked genes are strongly implicated in neuropsychiatric disease. We identified significant expression quantitative trait loci (eQTL) for 25,958 enhancers which mediate 6.8% of schizophrenia heritability, mostly independent from standard gene eQTL. Inclusion of enhancer eQTL in transcriptome-wide association studies enhanced functional interpretation of disease loci. Overall, our study characterizes the enhancer-gene regulome and genetic mechanisms in the human cortex in both healthy and disease states.

## Introduction

Enhancers are key regulatory regions of DNA that exert control over target gene expression from a distance^1,2^. Besides their critical function in orchestrating cell-lineage commitment and development^2,3^, enhancers play pivotal roles in mediating neuronal plasticity and memory formation in the brain^4^, and human evolved enhancers are hypothesized to drive advanced cognition^5–7^. Recent studies have found active enhancers are widely transcribed and enhancer expression levels represent an essential signature for enhancer activation^8–10^. Rather than merely being by-products of enhancer activity, several lines of evidence suggest that some, if not all, the enhancer expression products are functional^11–13^. More traditional approaches for identifying enhancers have their drawbacks: in contrast to transcribed enhancers (TEn), chromatin accessibility also captures non-active enhancers^14^ and histone modification marks, such as H3K27ac, can capture ‘‘stretch-enhancers’’ or ‘‘super-enhancers’’ of more than 10kb in size^15,16^, making it difficult to pinpoint functional loci. On the other hand, TEns are more likely to be validated in functional assays^10^, possibly due to the activity-dependent expression and smaller size. Although the Functional Annotation of the Mammalian Genome (FANTOM) project successfully annotated ∼65,000 TEns across more than 400 human tissues and cell types using only transcriptomic signatures captured by cap analysis of gene expression (CAGE-seq)^10^, it is thought that the majority of TEns have yet to be identified^8,9^. In the human brain, the annotation includes only a few thousand TEns and does not consider their distribution across the different cell types^10^. As such, a systematic cell-type-specific map of human brain enhancer expression would be an important step toward developing a more thorough understanding of enhancer functional units in the brain.

The majority of common neuropsychiatric disease-associated variants lie in the noncoding genome^17–21^, where brain-associated enhancers are overrepresented both for risk alleles and heritability of neuropsychiatric traits^22–26^. A plausible molecular mechanism is that noncoding variants affect enhancers and alter target gene expression. Indeed, studies based on epigenomic reference maps and large-scale gene expression quantitative traits loci (eQTL) have identified hundreds of genes and variants underlying disease risk, yet many more remain to be discovered^23,26–28^. Considering that enhancers outnumber expressed genes^29^, and multiple enhancers can regulate the same gene, eQTL analysis restricted to genes will likely miss critical information from genetic mechanisms mediated by enhancers under certain scenarios. Moreover, eQTLs are depleted near the critical genes for complex traits, including genes with redundant enhancers^30^, transcriptional factors, network hubs, and stress response genes^31^, motivating QTL analysis of enhancer function and their involvement in the etiology of neuropsychiatric disease. However, a population-level analysis of enhancers is hampered by limited access to biospecimens and available molecular assays. As enhancer expression levels reflect enhancer activity, it provides an alternative means to quantitatively investigate regulatory circuitry^10,32–36^. Although enhancer expression products are often not polyadenylated and, hence, are less stable, they can be captured by deeply sequencing total RNA^33,35,36^. Issues concerning the inherent instability and low expression rates of enhancers^37^ can be ameliorated when quantification is applied to large cohorts, allowing for the quantification of robustly expressed enhancers across many biological samples.

Here, we present a population-scale analysis of enhancer expression in the human cortex to understand the functional effect of genetic variations in neuropsychiatric disease. We generated comprehensive multi-omics reference maps for both neuronal and non-neuronal cells and used this to develop a computational scheme that accurately cataloged 30,795 neuronal and 23,265 non-neuronal TEns in the human cerebral cortex. We then examined the population-level variation of enhancer expression by analyzing 1382 RNA-seq libraries from 774 schizophrenia (SCZ) and control postmortem brains from the CommonMind Consortium (CMC). We explored enhancer-gene expression coordination and found that enhancer-linked genes are strongly implicated in neuropsychiatric diseases. We identified significant eQTL for 25,958 enhancers. The enhancer eQTLs are independent of gene eQTL, and mediate SCZ heritability complements standard gene eQT. Lastly, we performed transcriptome-wide association analysis (TWAS) of joint gene-enhancer eQTL, which greatly facilitated the functional characterization of schizophrenia risk loci.

## Results

### Comprehensive multi-omics maps of neuronal and non-neuronal cells

To annotate and characterize the expression pattern of TEns in the human cortex, we first performed multi-omics profiling, including ribosomal-RNA (rRNA) depleted total-RNA-seq, ATAC-seq, and ChIP-seq for H3K4me3 and H3K27ac) in neuronal (NeuN+) and non-neuronal (NeuN-) nuclei, isolated by fluorescence-activated nuclear sorting (FANS), from five brain regions (Brodmann areas 10, 17, 22, 36, and 44) of 10 control individuals (**Fig. 1a and 1b**). After extensive quality control, including assessing cell type, sex, and genotype concordance (see Suplementary Methods), a total of 14.5 billion uniquely mapped paired-end read pairs for RNA-seq (N=93), 4.3 billion for ATAC-seq (N=98), 11.0 billion for H3K4me3 ChIP-seq (N=96), and 9.6 billion for H3K27ac ChIP-seq (N=96) were obtained. Additionally, in a subset of samples (N=6), we mapped repressors and long-range chromatin interactions in neurons and non-neurons by performing H3K27me3 ChIP-seq and Hi-C, respectively.

**Fig. 1.**
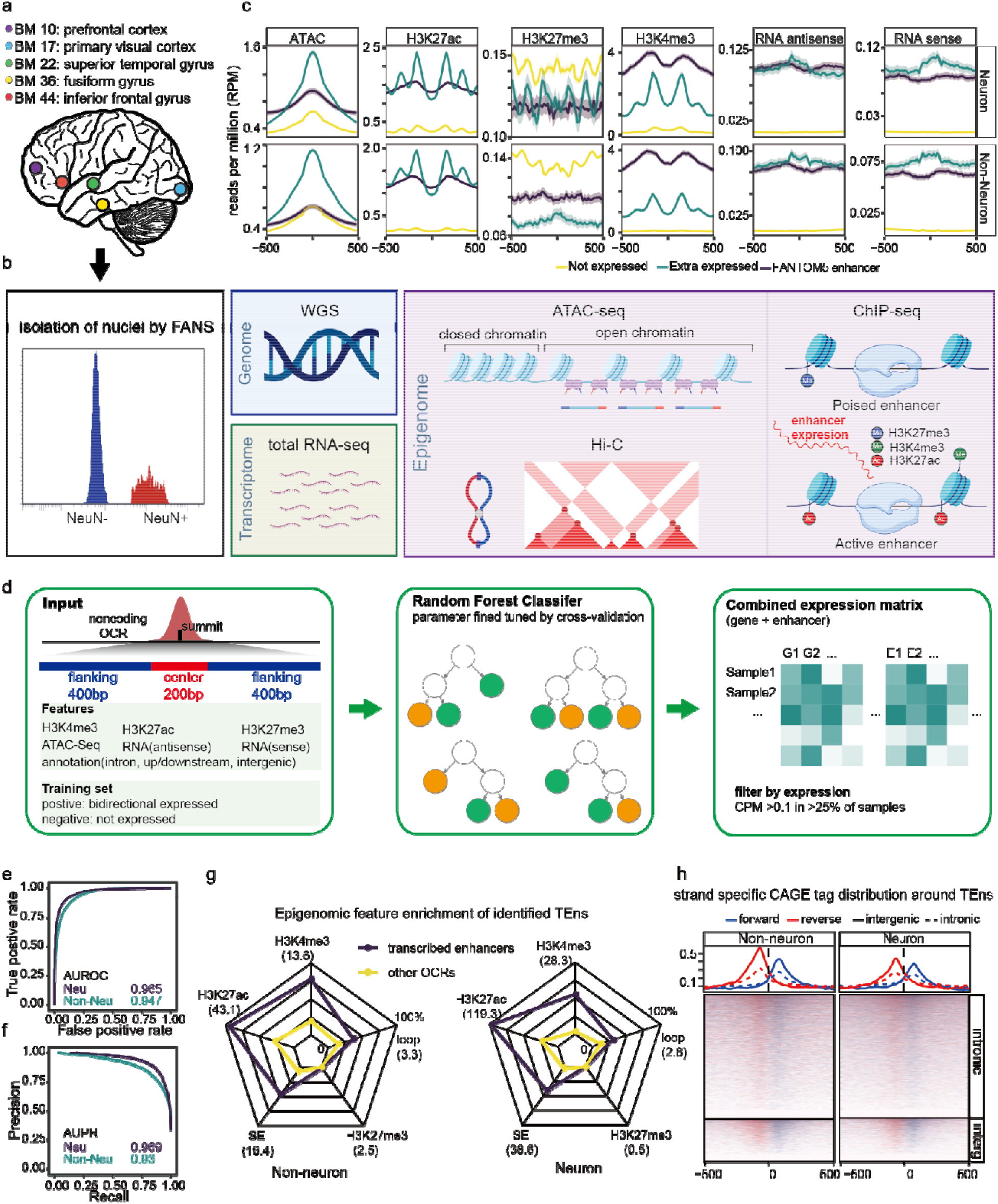
Catalog cell-type-specific TEns in the human cortex. **a**, Dissections from five brain regions (Brodmann area 10, 17, 22, 36, and 44) of 10 control subjects were obtained from frozen human postmortem tissue. **b**, Combined with FANS, we performed functional assays including total RNA-seq, ATAC-seq, H3K4me3/H3K27ac/H3K27me3 ChIP-seq, and Hi-C for neuronal (NeuN+) and non-neuronal (NeuN-) nuclei. **c**, Transcriptomic and Epigenomic profiles around expressed enhancers, including FANTOM 5 enhancers (FEs) and OCRs with bidirectional CAGE tags (Extra expressed, EEs), and non-expressed enhancers (NEs), the shadow shows the 95% confidence intervals. **d**, Flow charts demonstrate TEns identification pipelines. Briefly, transcriptomic and epigenomic signals around the enhancer regions were aggregated for both neuronal and non-neuronal cells. Expressed enhancers (FEs and EEs) were used as positive sets, and not expressed enhancers(NEs) were used as negative sets. The parameter was tuned with 10-fold cross-validation to select a random forest model to classify expressed and not expressed enhancers. Lastly, the enhancer expression matrix and gene expression matrix were combined for downstream analysis. **e**, ROC (true positive rate vs false positive rate) and **f**, PR (precision-recall) curve of the resulting random forest models exhibit high AUROC (area under the precision-recall curves) and AUPRC (area under the precision-recall curves). **g**, The radar plots show typical enhancer-related signals including H3K4me3, H3K27ac, H3K27me3, super-enhancer, and loop anchor occupancy between identified TEns and background (non-expressed enhancers). The value within parentheses indicates the odds ratio between identified TEns and background. **h**, Strand-specific CAGE tag average profile (top) and distribution (bottom) for intergenic and intronic TEns.

To validate the cell-type specificity of our data, we performed a transcriptome deconvolution analysis with reference markers from single-cell analysis^38^. The neuronal samples were strongly enriched for glutamatergic and GABAergic neurons, while the non-neuronal cells were enriched for oligodendrocytes, followed by astrocytes and microglia (**Extended Data Fig. 1a**), thus confirming the cell-type specificity of FANS. In addition, our ChIP-seq and ATAC-seq data were highly concordant with previously published reports (**Extended Data Fig. 1b**).

### A catalog of cell type-specific TEns in the human cortex

We sought to utilize our multi-omics dataset to further expand the catalog of TEns in the human brain. As an exploratory step, we compared the molecular profiles between expressed enhancers and those that were not expressed. To define a set of expressed enhancers, we collected FANTOM5 enhancers that overlapped with our H3K27ac peaks (FANTOM5 enhancers, FEs), as well as noncoding open chromatin regions (OCRs) that were missed by FANTOM5 but had bidirectional CAGE tags (Extra expressed, EEs). OCRs that did not overlap with FANTOM5 enhancers and had no CAGE tags were defined as non-expressed enhancers (Non-expressed, NEs) (**Fig. 1c**). We reasoned that the enhancer transcription initiation sites correspond to TF binding sites^8,9^, which can be determined by the ATAC-seq peak summit. Indeed, the FE positions, which were determined by CAGE tags, captured the ATAC-seq signal summit and were flanked by well-positioned nucleosomes. Its position relative to accessible chromatin and nucleosomes aligned well with the OCR-derived EEs and NEs, suggesting OCR summits pinpoint TEn positions (**Fig. 1c**). Both FEs and EEs exhibit local transcription signals, as well as typical active enhancer chromatin modifications, including H3K4me3 and H3K27ac enrichment (**Fig. 1c**). It’s worth noting that, compared to FEs, EEs had markedly lower levels of H3K4me3 signal, suggesting that FANTOM5 enhancers might be biased towards enhancers with high levels of H3K4me3. In contrast, NEs were depleted of such active enhancer histone marks and, in turn, displayed much lower expression levels.

Based on the distinct epigenomic signatures, as well as accurate enhancer positioning, we developed a supervised machine learning scheme to expand and capture the cell-type-specific TEns in the human brain (**Fig. 1d**). For each cell type, we used the central and flanking multi-omics signals, as well as the genomic annotation of the OCRs, as input for random forest models to select features differentiating expressed enhancers (positive sets, FEs, and EEs) from non-expressed enhancers (negative sets, NEs) (see Methods). The resulting models are of high accuracy, as measured by the area under the receiver operating characteristic (0.97 and 0.95 for neuron and non-neuron, respectively) and the area under the precision-recall curves (0.97 and 0.93 for neuron and non-neuron, respectively) (**Fig. 1e** and **1f**). Overall, we identified 36,927 neuronal and 27,379 non-neuronal TEns, among which only 2,487 (6.73%) of neuronal and 2,833 (10.3%) of non-neuronal TEns are within 2kb of any FANTOM5 enhancers. In sharp contrast to the background (i.e. non-expressed non-coding OCRs), the identified TEns exhibited strong active enhancer signatures (H3K4me3 and H3K27ac peaks) as well as long-range, chromatin interactions (**Fig. 1g**). Specifically, almost all (94% non-neuronal and 97.4% neuronal) TEns overlapped with H3K27ac peaks (odds ratio, OR=43.1 non-neuronal, OR=119.3 neuron, p<10^−16^ for both, two-sided Fisher’s exact test) and TEns were overrepresented in super-enhancer (SE) regions. On average, a single SE contained 7.47 and 6.50 TEns in neuronal and non-neuronal cells, respectively (**Extended Data Fig. 1e**) and, overall, 86% of neuronal and 91% of non-neuronal SEs had at least one TEn. In addition, TEns had a much higher density in SE regions (p<10^−16^ for both, two-sided Wilcoxon tests) (**Extended Data Fig. 1f**). To examine if the identified TEns represent distal transcription initiation sites, we superimposed strand-specific CAGE tags on the identified TEns. This approach provides a means to capture TSS, rather than elongation signals ^10,39^. As expected, both intergenic and intronic TEns exhibited bi-directional CAGE tags, confirming that the identified TEns represent bi-directional TSSs (**Fig. 1h**).

We then quantified the expression of both genes and TEns across neuronal and non-neuronal RNA-seq samples. We found that 30,795 (83.4%) neuronal and 23,265 (85.0%) non-neuronal TEns were expressed at >0.25 counts per million in >10% of the samples. We subsequently compared the differences in epigenome profiles between expressed TEns and expressed gene promoters (**Extended Data Fig. 1g**). As expected, TEns possessed a distinct epigenomic signature compared to typical promoters. Both protein-coding gene and long intergenic noncoding RNA (lincRNA) promoters displayed abundant H3K4me3 and H3K27ac signals, whereas TEns were enriched for H3K27ac but had much lower levels of H3K4me3. Moreover, TEns also had lower H3K27me3 levels compared to the protein-coding genes and lincRNAs.

### TEn expression captures cell-type-specific enhancer function

To further characterize neuronal and non-neuronal enhancers, we annotated chromatin states by jointly analyzing ChIP-seq data using ChromHMM (**Extended Data Fig. 2a and 2b**)^40^. Compared to other CREs, such as promoters and polycomb repressors, enhancers were markedly different between the two cell types (**Extended Data Fig. 2c**). In line with this, the identified TEns were also largely non-overlapping, confirming the strong cell-type specificity of enhancer elements (**Extended Data Fig. 2d**). We then quantified cell type differences by modeling the read count matrices of gene/TEns, and CREs from ATAC-seq and ChIP-seq, as well as confounders selected by covariate analysis (Methods and **Table S2**). Despite their modest expression levels, ∼90% of TEns were differentially expressed (DE) between the two cell types (**Extended Data Fig. 2e**). In addition, we assessed the variance explained by different factors (Methods), and cell type was the strongest source of variation for all assays (**Extended Data Fig. 2f**). The DE analysis identified 8,864 genes and 22,669 TEns that were upregulated in neurons, and 9,140 genes and 22,582 TEns upregulated in non-neurons. The effect size of cell-type differences between TEns expression and activities determined by ChIP-seq or ATAC-seq were highly correlated for both intergenic and intronic TEns (**Fig. 2a** and **Extended Data Fig. 2g**), highlighting that enhancer expression quantified by RNA-seq gives a good representation of enhancer activity.

**Fig. 2.**
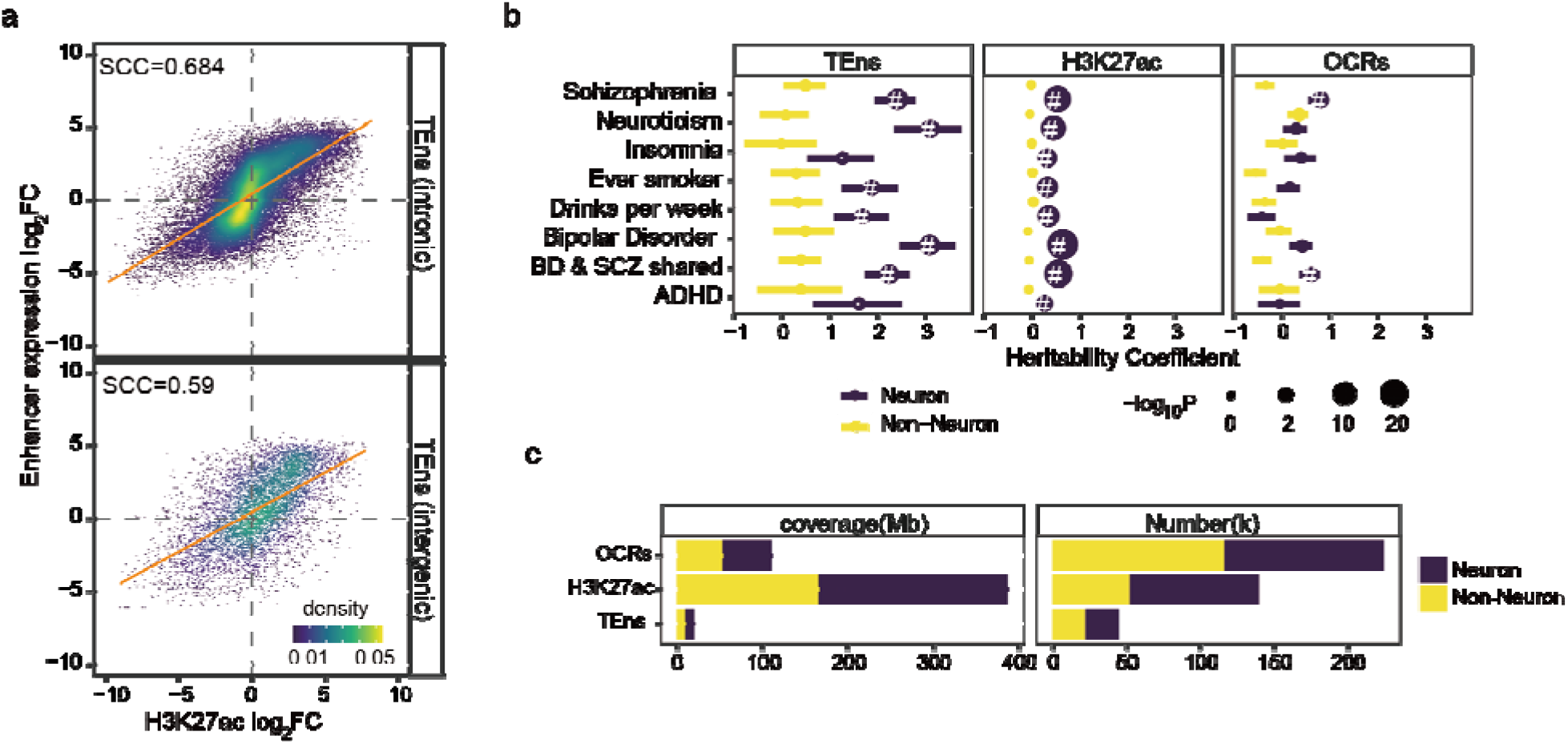
TEns expression captures cell-type-specific enhancer function. **a**, cell-type-specific effect size (log_2_ fold change) between TEns and overlapping H3K27ac peaks are highly consistent for both intergenic(N=7,624) and intronic(N=42,148) TEns. SCC represents the Spearman correlation coefficient (ρ) (p<10^−16^ for both). **b**, LD score regression enrichment for DE TEns/peaks of different neuropsychiatric traits. A positive coefficient signifies enrichment in heritability (per base enrichment). “·”: Nominally significant (p<0.05); “#”: significant after FDR (Benjamini & Hochberg) correction (FDR<0.05). **c**, DE TEns/peaks coverage (Mb) and numbers (k).

Given the strong colocalization of enhancer elements with common neuropsychiatric risk variants, we partitioned disease heritability with cell-type-specific TEns using linkage disequilibrium (LD) score regression^22^. In line with the result from H3K27ac peaks and OCRs, neuron-specific TEns were strongly enriched in risk variants for neuropsychiatric traits, including bipolar disorder (BD)^20^ and schizophrenia (SCZ)^41^. Conversely, enrichment in non-neuronal enhancers was not significant. Moreover, we examined the per-SNP heritability of these traits (Methods) and found that the per-SNP heritability of neuronal-specific TEns was remarkably higher than that of OCRs and H3K27ac peaks (**Fig. 2b**), suggesting a role for genetic regulation of cell-type-specific TEns in neuropsychiatric disease. Given the relatively small number and genomic coverage of TEns (**Fig. 2c**), TEns represent a smaller functional unit that accounts for a higher fraction of disease heritability.

### Enhancers are robustly expressed in independent cohorts

To systematically examine the population-level variation of enhancer expression in the human brain, in both healthy and disease states, we leveraged our cell type-specific brain TEns atlas to quantify the expression in a large-scale postmortem brain RNA-seq cohort collected and generated by the CommonMind Consortium^42,43^. A total of 1014 rRNA-depleted total RNA-Seq libraries, covering both the dorsolateral prefrontal cortex (DLPFC, BA9, and BA46) and anterior cingulate cortex (ACC, BA32, and BA24), as well as matched genotyping data from brain banks at the Icahn School of Medicine at Mount Sinai, the University of Pennsylvania and the University of Pittsburgh (a.k.a. MountSinai-Penn-Pitt, the CMC MPP cohort) were used as the discovery set (N_SCZ_ = 254, N_Control_ = 291, **Fig. 3a**). We utilized an independent, non-overlapping, cohort consisting of a total of 368 RNA-seq libraries, also derived from DLPFC and ACC, with matching genotype information from the NIMH HBCC brain bank (a.k.a. the CMC HBCC cohort) as a replicate cohort (N_SCZ_ = 78, N_Control_ = 151, **Fig. 3a**).

**Fig. 3.**
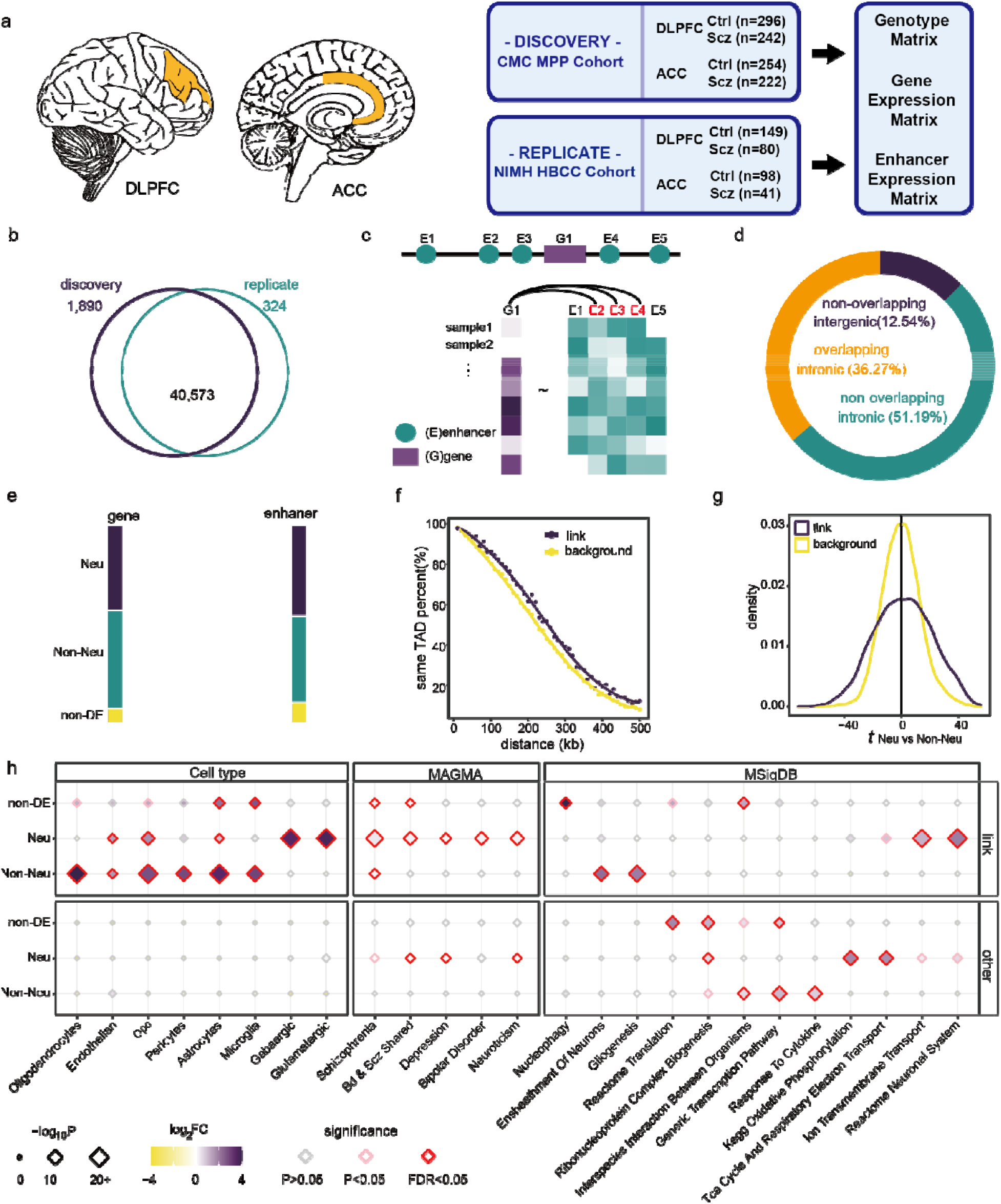
Enhancer-gene expression coordination. **a**, Two independent Cohorts were used for this study, both cohorts consisting of total RNA-seq from DLPFC and ACC postmortem brains as well as matched genotyping data. The discovery set includes 1,014 RNA-seq samples from N_SCZ_=254 and N_control_=291 control unique individuals. The replicate set includes 368 RNA-seq samples from N_SCZ_=78 and N_control_=151 unique individuals. Brain regions were visualized with cerebroViz^46^. **b**, The Venn diagram shows the overlap of expressed enhancers between the discovery and the replicate analysis. **c**, Demonstration of the lasso link model. For every gene, we considered all the enhancers within ±500kb of the TSS and fit a lasso model to select the linked enhancers. **d**, Distribution of the three different classes of enhancer-gene pairs: non-overlapping intergenic-enhancer gene pairs, overlapping intronic-enhancer gene pairs, and non-overlapping intronic-enhancer gene pairs. **e**, The differential expression status in the two cell types between gene and linked enhancers. **f**, Percentage of gene promoter and enhancer within the same TAD for different genomic distances at 10kb intervals. **g**, Compared to the background, the enhancer-linked genes have remarkably higher absolute t statistics between neuronal and non-neuronal cells (KS test, p<10^−16^). **h**, Cell types, neuropsychiatric common variants, and biological pathways enriched at different classes of linked and not-linked genes.

In the discovery cohort, we detected 42,463 enhancers and 20,704 Ensembl genes expressed at detectable levels (Methods). In the replication analysis, more than 95% of the enhancers detected in the discovery cohort were replicated (**Fig. 3b**, Jaccard index=0.948, p<10^−16^, two-sided Fisher’s exact test). In addition, the pairwise correlation between both intronic and intergenic enhancer expressions is highly consistent between cohorts (**Extended Data Fig. 3b**). Given the robust expression between the two independent cohorts, we combined the two data sets for the following analysis.

### *Cis*-coordination of expression between enhancers and target genes

Given that promoter-enhancer interactions can span considerable genomic distances, and not necessarily in a one-to-one manner, determining the target genes for enhancer elements is still a great challenge^1,3,44^. Our analysis enables a direct comparison of expression between enhancers and genes across multiple samples. To link enhancers to target genes, we took into account the joint effect of multiple enhancers, and fit a lasso regression model for every gene as a response variable and all enhancers within a ±500 Kb window (**Fig. 3c**). The resulting model detected 35,964 gene-enhancer links, consisting of 5,647 genes and 22,147 enhancers (**Table S4**). On average, a gene was linked to 5 enhancers (standard deviation, sd=5.34, **Extended Data Fig. 3c**), whereas an enhancer was linked to a single gene (sd=1.09, **Extended Data Fig. 3d**), in agreement with a previous estimation^45^. The majority of the associations are between non-physically-overlapping gene-enhancer pairs (**Fig. 3d**). Consistent with the positive regulation of active enhancers, the majority of the enhancer links (75.98%) had positive coefficients. As expected, enhancer-linked genes had a significantly higher expression level relative to the background (expressed genes without linked enhancer) (**Extended Data Fig. 3c**, p<10^−16^, two-sided Wilcoxon test). Moreover, enhancers originating from SEs were more likely to be coordinated with gene expression (p<10^−16^, OR=1.35, two-sided Fisher’s exact test); more than 65% of the linked enhancers come from SE loci, highlighting the critical role of SEs in gene regulation. Although we included all enhancers within a ±500 Kb window to the TSS, the significant enhancers were located closer to the TSS (p<10^−16^, two-sided Wilcoxon test compared to background, **Extended Data Fig. 3e**), even after excluding the physically-overlapping gene-enhancer pairs from the analysis (p<10^−16^, two-sided Wilcoxon test compared to background, **Extended Data Fig. 3f**). In addition, the cell-type specificity between linked pairs was also highly concordant (**Fig. 3e**), indicating that our model captures cell-type-specific regulatory mechanisms.

An alternative way of determining promoter-enhancer interactions is by directly measuring the physical contact of DNA using chromosome conformation capture technologies, such as Hi-C. These approaches have found two prominent chromatin structural features that are associated with enhancer-promoter interactions; long-range chromatin loops; and topological associated domains (TAD), which insulate the interactions across domain boundaries^47^. Compared with the background, the enhancer-gene-linked pairs had a significantly higher chance of being linked by chromatin loops (OR=1.84, p<10^−16^, two-sided Fisher’s exact test). In addition, most of the identified promoter-enhancer pairs were within the same TAD (59.63%, OR=1.81, p<10^−16^, two-sided Fisher’s exact test). We found that the linked enhancer-gene pairs are enriched for being within the same TAD compared to regions an equivalent distance apart (**Fig. 3f**). Taken together, our gene-enhancer links are well supported by chromatin interaction features, and our approach established cell-type-specific regulatory mechanisms linking enhancers to their target genes.

### Enhancer-linked genes are implicated in neuropsychiatric disease

Given that enhancers drive cell-type-specific gene expression, we sought to determine whether enhancer-linked genes are more likely to be cell-type-specific. By examining the *t*-statistics from differential expression (DE) analysis between between neuronal and non-neuronal cells, we found that enhancer-linked genes are strongly DE between the two cell types, compared to expressed genes that are not linked (p<10^−16^, Kolmogorov–Smirnov (KS) test, **Fig. 3g**). To dissect the function of enhancer-linked genes, we annotated the genes into neuronal and non-neuronal categories based on the DE analysis. As expected, enhancer-linked genes were strongly enriched for the main CNS cell types annotated from single-cell analysis^38^. The linked neuronal genes were strongly enriched for glutamatergic and GABAergic neurons, and non-neuronal genes were enriched for glial cell types such as oligodendrocytes and astrocytes (**Fig. 3h**). In contrast, non-linked genes were not even nominally significant in any of the cell types.

Previous studies have highlighted the function of neuronal genes in the etiology of SCZ^48^. For neuronal genes we found that both enhancer-linked and non-linked genes were strongly enriched for common neuropsychiatric trait variants, and that the enrichment level was remarkably higher in enhancer-linked genes, especially for SCZ (**Fig. 3h**). Aside from the neuronal genes, enhancer-linked non-neuronal and non-DE genes were also enriched for the SCZ GWAS signal (**Fig. 3h**), indicating that our model captured SCZ functional genes across different cell types. In addition, the neuronal enhancer-linked genes were enriched for SCZ related pathways, including ion transmembrane transport and the Reactome neuronal system^49^; the non-neuronal enhancer-linked genes were enriched for known pathways that are also implicated in SCZ, including ensheathment of neurons and gliogenesis^50^. Together, we found that enhancer-linked genes were strongly implicated in SCZ, highlighting the critical role of enhancers in cell-type-specific functions in the CNS and neuropsychiatric disease.

### Molecular QTL analysis highlights cell-type-specific enhancer regulation

To explore the genetic regulation of enhancers, we first examined the expression *cis*-heritability, which measures the fraction of expression variance explained by SNPs within the *cis*-window. Although lower than that of genes, we observed a substantial proportion of enhancers are *cis*-heritable (p<0.05) (**Extended Data Fig. 4a**). In addition, the heritability of enhancers was reproducible between the two brain regions (DLPFC and ACC; OR=7.29, p<10^−16^, two-sided Fisher’s exact test, **Extended Data Fig. 4c**).

We then performed joint eQTL analysis for both genes (GeQTL) and enhancers (EeQTL). For DLPFC and ACC brain regions, gene-enhancer combined expression matrices of European ancestry were independently adjusted for both known and surrogate covariates. An eQTL meta-analysis was subsequently performed with the normalized expression matrices across two brain regions using a linear mixed model to maximize power and account for repeated measures^51^. The model identified 3,593,102 *cis*-eQTL variant at FDR ≤ 5%, including 1,001,939 EeQTLs regulating 25,958 (62.86% of autosomal) enhancers, as well as 2,591,163 GeQTLs regulating 16,165 (86.46 % of autosomal) genes (**Table S5**). The most significant SNP (eSNP) for both genes (GeSNP) and enhancers (EeSNP) was centered around the corresponding transcription start site (**Extended Data Fig. 4e**). In line with previous findings that QTLs jointly influence multiple molecular phenotypes^52^, more than half (56.29%) of the EeQTL encompassed SNPs are also GeQTLs. However, eSNPs were largely independent between genes and enhancers. Only 6.1% of the EeSNPs reside within any GeSNP LD blocks (r^2^>0.5) (**Fig. 4b**), and only 12.9% of GeSNPs were located in the EeSNP LD blocks (r^2^>0.5) (**Fig. 4c**), highlighting that EeQTLs represent distinct genetic regulation information.

**Fig. 4.**
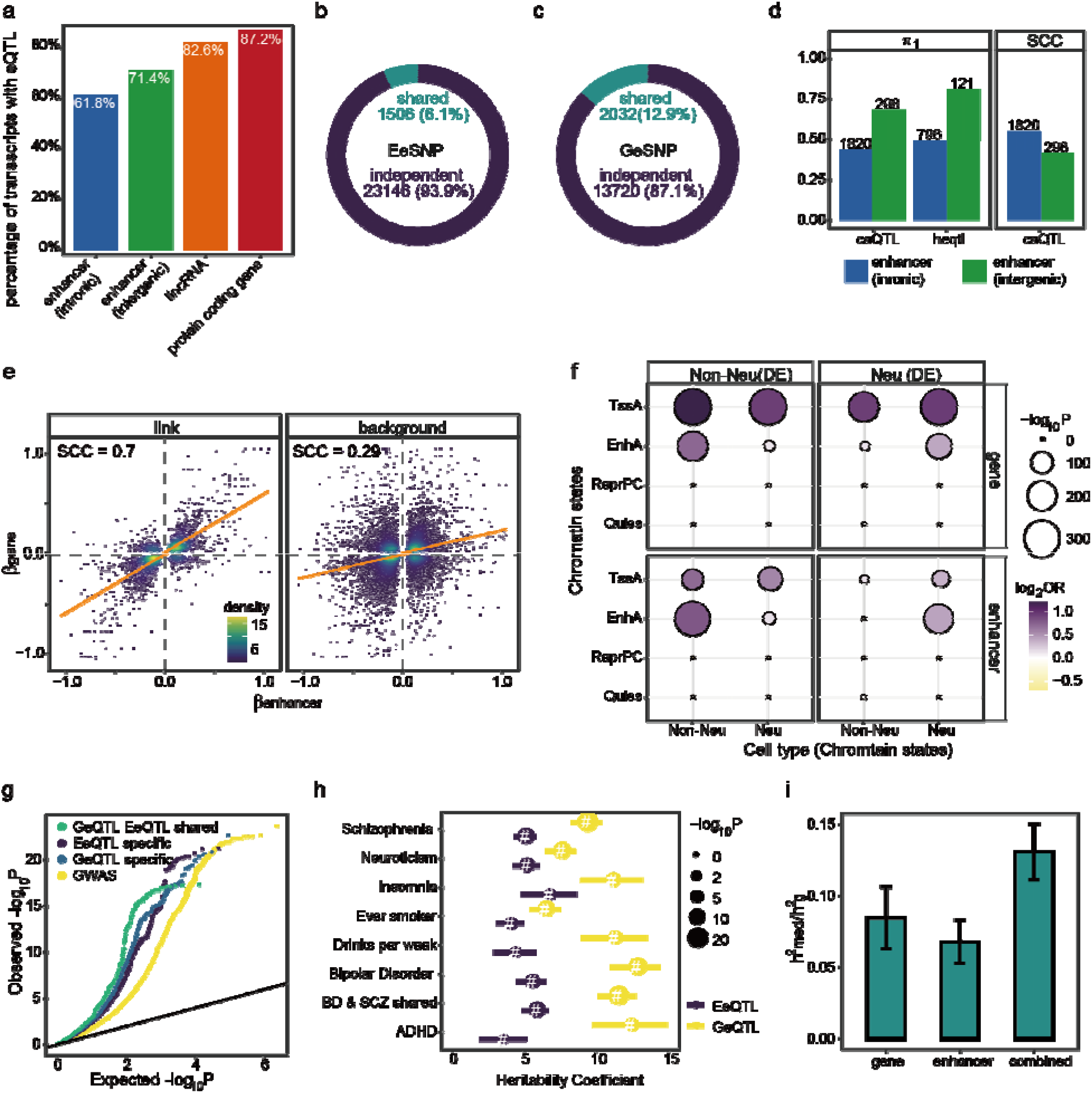
Genetic effects of enhancer expression. **a**, The percentage of different classes of autosomal gene/enhancers that have significant eQTLs. **b**, Percentage of EeSNP resides within any GeSNP LD blocks (r^2^>0.5). **c**, Percentage of GeSNP resides within any EeSNP LD blocks (r^2^>0.5). **d**, The replication of reported hQTL and caQTL in our analysis. Storey’s Π_1_ values for significant hQTL and caQTL in the EeQTLs. SCC values are the Spearman correlations (ρ) of effect sizes between caQTL and corresponding EeQTL. The size of the point corresponds to the number of unique enhancers used. The number above the bar corresponds to the count of unique enhancers used. **e**, The allelic genetic effect between gene and target enhancers are highly consistent. **f**, Enrichment of neuronal and non-neuronal genes and enhancers corresponding eQTLs in four chromatin states: TssA (active promoter), EnhA (active enhancer), ReprPC (polycomb repression), and Quies (other). The filled color represents enrichment fold change, and the size corresponds to enrichment P values. **g**, Quantile–quantile plot of SCZ GWAS p values. EeQTL specific, eQTL specific, and shared SNP are shown in comparison with genome-wide SNPs. GWAS SNPs were binarily annotated using SNPs within r^2^ >0.8 of the eSNP. **h**, LD score regression enrichment for gene and enhancer eSNPs of different neuropsychiatric traits. Positive coefficient signifies enrichment in heritability (per base enrichment).”·”: Nominally significant (p<0.05); “#”: significant after FDR (Benjamini & Hochberg) correction (FDR<0.05). **i**, Estimated proportion (± standard error) of heritability mediated by the cis genetic component of assayed enhancer, gene, and combined expression for SCZ.

To confirm our eQTL result, we first compared our GeQTL with the GTEx brain tissues^53^ and observed high Π_1_ values (median 0.950, **Extended Data Fig. 4f**) as well as the high concordance of allelic effect (**Extended Data Fig. 4f**). We then compared the EeQTL with published brain histone QTL (hQTL)^54^ and chromatin accessibility QTL (caQTL)^55^ data. We observed high Π_1_ values for EeQTLs (**Fig. 4d**). Moreover, both intergenic and intronic EeQTLs exhibited strong positive correlations comparing the effect sizes with caQTL (**Fig. 4d**), indicating that EeQTL captures enhancer genetic regulation. We next examined the genetic effect between the linked enhancer and gene pairs. We reasoned that genetic variants would have a concordant effect on functional gene-enhancer pairs. Indeed, we observed the effect sizes were highly consistent between linked genes and enhancers (**Fig. 4e**) and were independent of the overlapped gene-enhancer effect (**Extended Data Fig. 4g**).

Having shown that TEns play pivotal roles in cell-type-specific gene expression, we next sought to test if genetic regulation of TNs drives cell-type-specific gene expression. Under this model the cell-type-specific gene eQTL will be enriched in corresponding enhancer regions. To test this, we grouped GeSNPs based on the DE status between neuronal and non-neuronal cells and annotated the SNPs with cell-type-specific chromatin states using GREGOR^56^. As expected, gene eSNPs were strongly enriched for known CREs, including active promoters (TssA) and active enhancers (EnhA) (**Fig. 4f**). As distinct from that of the promoters, the enrichment of gene eSNPs at enhancers was highly cell type-specific, highlighting the central role of enhancer elements in cell-type-specific genetic regulation. In contrast to GeSNPs, EeSNPs were strongly enriched in enhancers of corresponding cell types instead of active promoters, distinguishing EeQTLs from GeQTLs.

### EeQTL mediate schizophrenia heritability

As described above, neuronal-specific enhancers are enriched for SCZ GWAS signals (**Fig. 2c**). In addition, we found that both EeQTL unique SNPs, and those that are shared with GeQTL SNPs, exhibit an excess of low P values for the SCZ GWAS signal (**Fig. 4g**). Taken together, these findings indicate that EeQTLs contribute to SCZ heritability in a complementary manner to GeQTLs. Given the relative independence of eSNP signals, we subsequently performed LD score regression to quantify the GWAS signal enrichment at both EeSNP and GeSNP loci. As expected, we found both EeSNPs and GeSNPs are strongly overrepresented with SCZ common variants as well as other neuropsychiatric traits (**Fig. 4h**).

Given that both GeQTLs and EeQTLs are associated with SCZ heritability, we next asked if the heritability is mediated by gene and enhancer eQTL, or if it arises due to other non-causal situations such as linkage and/or pleiotropy. We estimated the proportion of SCZ heritability mediated by gene/enhancer expression (h^2^_med_/h^2^_g_) using MESC^57^, which quantifies the proportion of disease risk mediated by QTLs. A substantial fraction (6.8 1.5%) of SCZ heritability is mediated by enhancer expression (**Fig. 4i**), indicating that enhancer is a key causal factor in the etiology of SCZ. In addition, we combined the gene and enhancer expression matrix and found that the mediated heritability proportion increased from 8.4 2.2% (gene) to 13 1.9% (combined), suggesting that genes and enhancers mediate their genetic effects complements standard gene eQTL.

### Adding enhancers to transcriptome-wide association studies facilitates fine-mapping and interpretation of SCZ GWAS loci

Having shown that SCZ genetic variants contribute substantially to gene and enhancer expression, we next sought to identify the potential genes and enhancers that mediate the genetic risk for SCZ. We performed transcriptome-wide association studies (TWAS)^58^ using gene and enhancer eQTLs, and the most recent SCZ GWAS^41^. Initially, we built elastic net and lasso regression genetic variant-based expression prediction models^59^ for *cis*-heritable transcripts in DLPFC and ACC brain regions. This resulted in expression models for 10,669 unique genes and 8,702 unique enhancers from the two brain regions, which markedly increased transcriptome coverage for TWAS compared to previous human brain studies in SCZ^28,60–62^. The models yielded 204 genes and 98 enhancers that are outside of the MHC region and are associated with SCZ (Bonferroni-adjusted P, P_bonferroni_<0.05), covering 104 of 264 non-MHC autosomal independent genome-wide significant SCZ GWAS loci (**Fig. 5a**). Specifically, 26 loci are shared between enhancers and genes, 23 loci are only tagged by enhancers and for 55 loci only genes were detected. To confirm the reproducibility of our studies, we compared the TWAS *Z* scores between the two brain regions, as well as the *Z* scores between our gene models and previous reports^28,62^, where both comparisons exhibit high concordance (**Extended Data Fig. 5a** and **5b**).

**Fig. 5.**
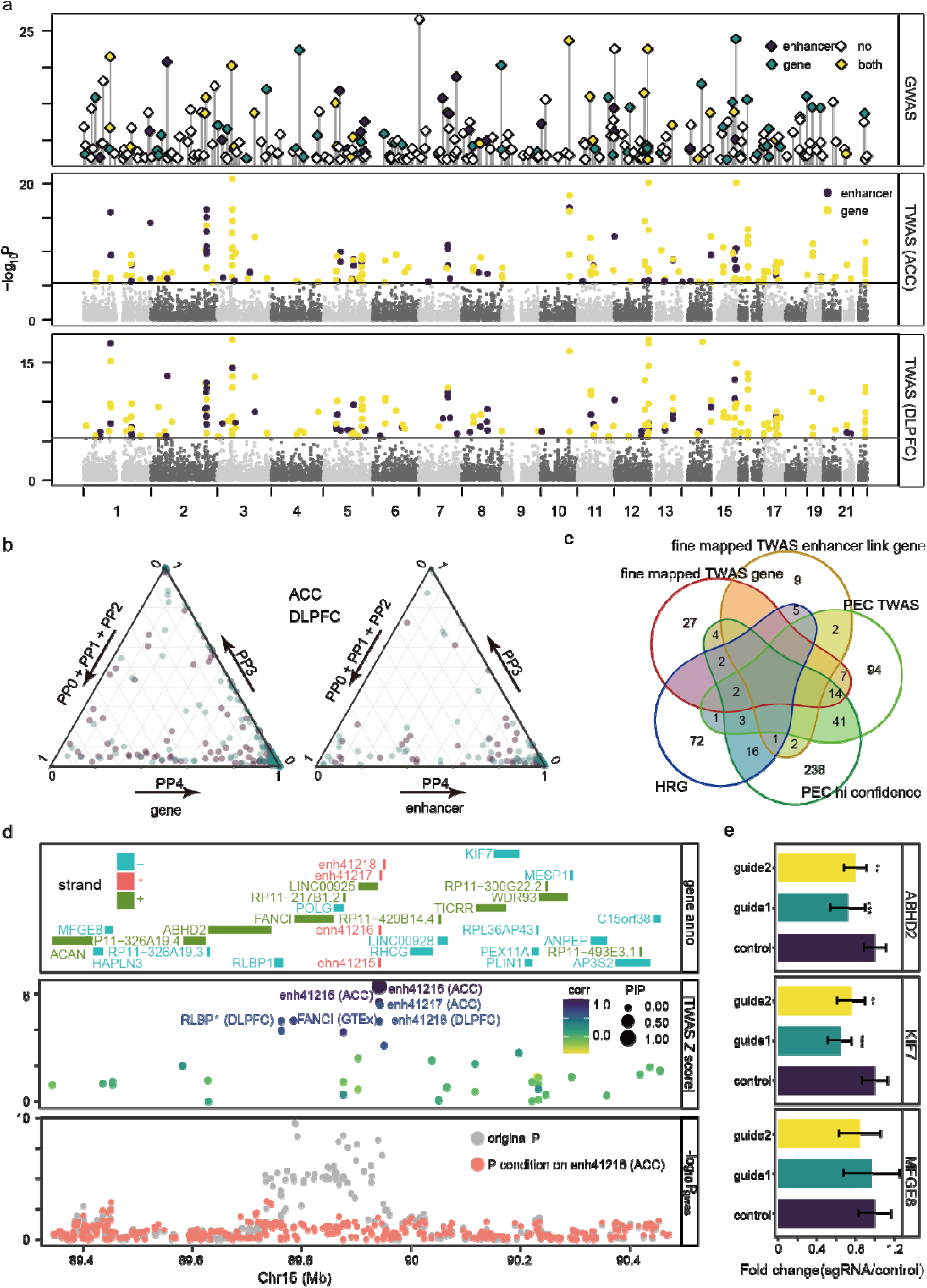
SCZ TWAS. **a**, Manhattan plot of SCZ TWAS enrichment in DLPFC, ACC, and the independent genome-wide significant SCZ associations (excluding chrX and MHC). GWAS node height corresponds to the index SNP significance, and the color indicates if the GWAS loci are associated with enhancer, gene, both gene, and enhancer, or none of them. Significant TWAS enhancers (purple) and genes (yellow) are highlighted in different colors. **b**, Ternary plots showing coloc posterior probabilities for significant TWAS genes and enhancers respectively. PP0+PP1+PP2: three scenarios for lack of test power; PP3: independent causal variants; PP4: colocalized causal variants. **c**, Venn diagrams show the overlap between the fine mapped TWAS genes, TWAS-enhancers linked genes, PsychENCODE (PEC) TWAS genes^28^, PsychENCODE(PEC) high confidence risk genes^23^, and SCZ high-confidence risk genes (HRG)^67^. **d**, Illustration of genomic loci at chr15, harboring multiple TWAS loci. Top panel, transcript position (only cis-heritable transcripts are shown), the color indicates transcription direction of genes. Middle, gene/enhancer TWAS Z score absolute value, point size indicates the FOCUS posterior inclusion probability (PIP), color indicates the genetic correlation with the highest PIP. Bottom, Manhattan plot of SCZ GWAS signal, before and after conditioning on enh41216. **e**, KRAB-dCas9-mediated repression of enh41216 leads to the reduction of ABHD2 and KIF7 expression as measured by qPCR in neural progenitor cells. (^***^ p<0.001, ^**^ p< 0.01, ANOVA, N=6, mean ± standard error are shown)

To avoid spurious associations from LD structure^63^, we first checked if the eQTL and GWAS association is driven by the same causal variants. We conducted a colocalization analysis^64,65^ to estimate the probability that the eQTL and GWAS signals are associated (PP4) or not (PP0, PP1, PP2 and, PP3). We found both genes and enhancers exhibited high PP4 value (**Fig. 5b**), 66 of the 98 TWAS enhancers, and 139 of the 202 TWAS genes colocalized between eQTL and GWAS signals (PP4>0.5). We next conducted a fine-mapping analysis that controls for the correlation structure introduced by LD and SNP weights, as well as certain pleiotropic effects to refine the TWAS associations using FOCUS^66^. To account for genes that are filtered by low *cis*-heritability, we included gene prediction results from GTEx expression panels. The analysis yielded a credible set of 466 transcripts, including 384 genes and 82 enhancers covering 151 GWAS loci, where 20 of the loci were only tagged by enhancers and 29 of the loci were tagged by both gene and enhancers.

To further refine the transcript-based fine-mapping of the TWAS outcome, we required the genes and enhancers to (i) have P_bonferroni_ < 0.05 in the TWAS analysis, (ii) have PP4 > 0.8 in the colocalization analysis, and (iii) be within the credible sets of FOCUS transcripts, resulting in 16 enhancers and 56 genes (**Table S6**). Although a substantial percentage of the associated genes have been identified previously, many of them are novel (**Fig. 5c**). Using our gene-enhancer link model, we identified 19 genes (MAN2A1, VRK2, FANCL, LINC01877, FTCDNL1, DPYD, CTNND1, MFGE8, ABHD2, STH, SATB2, C2orf69, MAIP1, PJA2, FAM114A2, MMP16, KIF7, AC092691.1, SAP30L) that are associated with the fine mapped enhancers (**Fig. 5c** and **Table S6**), many of which have been reported previously^23,28,67^ (**Fig. 5c**) but have not been directly identified by TWAS analysis.

We provide an illustrative example for an SCZ locus that includes enh41216 (**Fig. 5d**), a novel TEn that resides within a SE. The GWAS locus is associated with 12 TEns and 32 genes but only enh41216 is identified as a causal transcript by FOCUS fine-mapping. The top SNP of this enhancer, rs2247233, is also one of the top GWAS SNP (**Extended Data Fig. 5c**). In addition, we performed a conditional analysis and found that the enh41216 TWAS signal fully explained the GWAS significance (**Fig. 5d**, bottom).

The enhancer is linked to three genes, ABHD2, MFGE8, and KIF7. To experimentally validate the enhancer target, we performed CRISPR interference of enh41216 in neural progenitor cells (**Fig. 5d**). This led to a 20-30% reduction in gene expression for KIF7 and ABHD2, two genes that reside ∼200kb from the enhancer, whereas levels of MFGE8, 450kb from the enhancer, remained unchanged. KIF7 has been known to play a role in regulating neuronal development and brain abnormalities in ciliopathies^68^, but was not significant in the TWAS analysis (TWAS P_bonferroni_>0.1, FOCUS posterior inclusion probability, PIP<0.001). Moreover, knockdown of KIF7 results in thinner and shorter processes of multipolar neurons^69^. ABHD2, which is not significant in the TWAS analysis (TWAS P>0.1, FOCUS PIP<0.001), encodes an enzyme that catalyzes the hydrolysis of 2-arachidonoyl glycerol, which is a signaling lipid in the CNS and a key regulator of neurotransmitter release^70,71^.

## Discussion

A growing body of evidence suggests that enhancer elements play a pivotal role in neuropsychiatric diseases^19,22,23^. However, a systematic view, including regulatory circuits, and genetic effects of enhancer sequences is lacking. Here, we utilized population-level variation of gene and enhancer expression in the human brain to provide a comprehensive assessment of the regulatory mechanisms of transcribed enhancers in SCZ. We illustrate how to leverage large-scale transcriptome data to investigate enhancer regulatory circuits and provide novel insights into complex traits.

Given that enhancers are modestly expressed, previous attempts to identify TEns using a single assay either used a stringent threshold and missed the majority of TEns^10^, or, with a more flexible cutoff, identified mostly non-enhancer elements^72^. In our analysis, we used a two step approach to define brain TEns with increased sensitivity and specificity. We initially performed cell-type-specific TEn discovery based on supervised machine-learning integrative analysis of multi-omics data. We then further refined TEn mapping and studied the properties of the enhancer regulome using population-level expression from RNA-seq data. This approach is very sensitive, resulting in 30,795 neuronal and 23,265 non-neuronal TEns, expanding, by an order of magnitude, the repertoire of known transcribed enhancers in the human brain. In terms of specificity, the accuracy of our method is confirmed by active enhancer histone modification occupancy (∼95% H3K27ac), bidirectional transcription initiation signal, as well as the high precision-recall of our predictive model. Compared to the traditional and broad active enhancer assays, H3K27ac and H3K4me1 ChIP-seq, our TEns-based approach has the advantage of higher resolution to define smaller functional regions. In addition, our approach can better refine the chromatin accessibility landscape by subsetting ATAC-seq peaks that represent active enhancers from other regulatory sequences. In line with this, analysis of cell type-specific TEns identifies markedly higher SNP heritability for neuropsychiatric traits than regulatory sequences defined using ATAC-seq and H3K27ac ChIP-seq.

Based on the coordinated expression of enhancers and target genes, we utilized population-level variation of the transcriptome and regulome in 1,382 brain samples to model *cis*-coordination and determine gene-enhancer interactions. As enhancers can exert their effects across long genomic distances, the closest gene is not necessarily the target gene. Spatial interactions determined by Hi-C are limited by the resolution of the method and by bias due to proximity ligations^73^, and may have other functions besides facilitating promoter-enhancer interactions. Our approach is validated by the concordance with chromatin organization features, consistent allelic genetic effects, and *in vitro* CRISPR inference validation. Enhancer-linked genes are highly expressed, are strongly enriched for CNS cell types and neuropsychiatric trait risk loci, highlighting the importance of enhancers in the etiology of disease.

Given the significant *cis*-heritability of enhancer expression, we performed eQTL analysis to link *cis* genetic variation to enhancer expression^51^. We found that EeQTLs were (i) highly concordant hQTLs and caQTLs, (ii) centered around the enhancer loci, (iii) relatively independent of GeQTLs, and (iv) enriched in active enhancer chromatin states, suggesting that enhancer expression represent enhancer activity instead of transcriptional noise. Both TEns and EeSNPs confer substantial neuropsychiatric disease liability. We subsequently determined the proportion of SCZ heritability mediated in *cis* by enhancer expression^57^ and found that enhancer expression alone, or combined with gene expression, explained a substantial percentage of SCZ *cis*-genetic variants (6.8% and 13%, respectively). We performed TWAS and fine-mapping and found that a significant fraction of the SCZ GWAS loci is captured only by enhancers. enh41216 was prioritized based on fine-mapping in an SCZ locus containing 12 TEns and 32 genes. enh41216 was predicted to regulate three genes, among which two (*KIF7* and *ABHD2*), located ∼200kb from the enhancer, were validated using CRISPR interference in neural progenitor cells. Taken together, our study further emphasizes the importance of the regulome as an additional layer to functionally characterize disease vulnerability.

Overall, the consideration of population variation in enhancer expression in large-scale total RNA-seq analysis provides novel insights into the regulatory mechanisms of gene expression, as well as the genetic effects influencing complex traits. The framework described in this study provides an inexpensive and high-resolution approach to explore active enhancer function in most human tissue and cell types.

## Methods

### Multi-omics human postmortem brain samples

Brain specimens from 10 neurotypical individuals were obtained from the Mount Sinai/JJ Peters VA Medical Center Brain Bank (MSBB–Mount Sinai NIH Neurobiobank), as part of the Accelerating Medicines Partnership - Alzheimer’s Disease (AMP-AD) project^74^. All neuropsychological, diagnostic, and autopsy protocols were approved by the Mount Sinai and JJ Peters VA Medical Center Institutional Review Boards. Details about the subject information, including sex, age, postmortem interval (PMI) could be accessed on Synapse via the AD Knowledge Portal (https://adknowledgeportal.org).

The details of the experimental procedural, data processing, and quality control information of the molecular assays can be found in the supplementary material.

### CRISPR inference validation

Cell culture and transduction of dCas9-effector hiPSC-NPCs with gRNA lentivirus. The iPSC lines used in this study (NSB2607-1-4 and NSB553-S1-1)^75,76^ were maintained in Matrigel (Corning, cat.# 354230) coated 6-well plates under NPC medium (DMEM/F12 (Life Technologies, cat.# 10565), 1x N2 (Life Technologies, cat.# 17502-048), 1x B27-RA (Life Technologies, cat.# 12587-010), 20 ng/ml FGF2 (R&D Systems, cat.# 233-FB-01M). Upon confluency, cells were dissociated with Accutase (Innovative Cell Technologies, cat.# AT104) for 5 minutes at 37°C, quenched with DMEM/F12, pelleted, and resuspended in NPC medium containing 10µM/ml Thiazovivin (THX) (Sigma/Millipore, cat.# 420220). 3.5×10^5 NPCs per well were seeded onto Matrigel-coated 24-well plates in NPC media. The following day, gRNA lentiviruses such as LentiGuide-Hygro-mTagBFP2 (Addgene, cat.# 99374) were added to cultures, followed by spinfection (1 hour, 1000xg, 25°C). After spinfection, the cultures were incubated overnight, and medium was then replaced the following day. The cells were selected with 0.3 μg/ml puromycin for dCas9-KRAB (Sigma, cat.# P7255) for two days. After puromycin selection, cells were fed with fresh NPC medium and then harvested two days later. gRNA expression was confirmed via BFP fluorescence, prior to harvest.

The Benchling CRISPR gRNA design tool was used to design guide sequences. Two guide RNAs for enh41216 with the highest specificity scores were selected:

Guide RNA1 (chr15: 89400303): 5’-GAGGCGCGATACGAACCCGT-3’

Guide RNA2 (chr15: 89400387): 5’-GATACGGGCGAATCCCGCAA-3’

The guide oligos were then synthesized by IDT (Integrated DNA technologies) and phospho annealed via T4 PNK (37°C for 30min, 95C for 5min and ramped down to 25°C at 5°C/min). The annealed oligos were then cloned downstream of a constitutively expressed U6 promoter in a lentiviral vector (lentiGuide-Hygro-mTagBFP2, Addgene, cat.# 99374) using the golden gate cloning method (Bsmb1; 30 cycles of 37°C for 5min and 20°C for 5min). Vectors were packaged into 3rd generation lenti-viruses by VectorBuilder and transduced into dCas9-KRAB expressing neural progenitor cells (NPCs). RNA was extracted from transduced NPCs and target gene expression knockdown was validated through TaqMan RT-qPCR relative to cells that were transduced with a negative control scrambled guide. The 2–ΔΔCt method was used to determine fold change in expression relative to the housekeeping genes GAPDH and ACTB.

### TEns identification

Training set. To define a training set, we first collected the brain-associated FANTOM5 enhancers that overlapped with our H3K27ac peaks (FANTOM5 enhancers, FEs). Given that FANTOM5 enhancers only cover a small fraction of TEns, and could have bias, we next determined the CAGE tags at the non-coding OCRs with R bamsignals package (v1.14.0) in a strandspeicifc mode. The analysis yielded 6,810 neuronal and 5,509 non-neuronal bidirectionally expressed enhancers (EEs), as well as 97,827 neuronal and 45,387 non-neuronal not expressed enhancers (NEs). We subsequently compared the epigenomic and transcriptomic profiles at the three sets of enhancers with the bamsignals package (https://bioconductor.org/packages/bamsignals), confirming that the two expressed enhancer groups exhibit different features compared to the not expressed enhancers. We only used the FEs that overlapped corresponding OCRs for downstream analysis. We used FEs and subsampled the same amount of EEs as the positive set. A comparable number of negative sets is subsampled from the NEs.

#### Testing set

To define a test set, we focused on ATAC-seq peaks. Peaks that overlapped with an annotated exon as well as a 1 kb region up/downstream (Gencode V30)^77^, ribosome DNA loci^78^, and ENCODE blacklisted regions^79^ were filtered. We extend the resulting peak summit to 500bp based on the distribution of FANTOM5 enhancer size (**Extended Data Fig. 1c**), resulting in 168,841 neuronal and 93,139 non-neuronal enhancers.

#### Prediction model

We collected tag counts with ATAC-seq, ChIP-seq, and strand-specific RNA-seq bam files for the central 200bp and flanking 400bp of both the testing and training sets (**Fig. 1d**). Additionally, we annotated the position relative to genes. The BAM counts and genomic annotation of the training set were used as the input for a random forest model^80^, with parameters fine-tuned by a 10-fold cross-validation grid search^81^. The performance of the resulting models was determined by AUROC and AUPC^82^, both of which generate values of ∼0.95 (**Fig. 1e and 1f**). We subsequently predict the TEns from testing sets with the trained models for both cell types.

#### Quantification

We counted the number of reads overlapped with the TEns using featureCounts function in RSubread (v1.6.3)^83^. The enhancer-gene combined expression matrix (counts per million, CPM) was used for the following analysis.

### Gene/TEn/peaks filtering and Differential analysis

We performed differential analysis between neuronal and non-neuronal cells for genes/enhancers with RNA-seq, and peaks of ATACseq and ChIP-seq from the Multi-omics cohort. For RNA-seq analysis only protein-coding genes, lincRNA, and TEns were used. A consensus read count-based differential analysis pipeline was used with the following steps:

#### Read count and transcript/peak filtering

For each assay, transcript/peak read count matrices were used as input. For ATAC-seq, H3K4me3, and H3K27ac ChIP-seq, only peaks that had at least 1 count per million reads (CPM) in > 10% of the samples were retained. For Multi-omics RNA-seq data, we used a relatively low cut-off to account for the modest expression of TEns and retained all the transcripts that had at least 0.25 CPM in >10% of the samples. The retained TEns that overlapped between neurons and non-neurons were merged and used for downstream analysis. For the CMC data, we contained all the transcripts that had at least 0.1 CPM in >40% of the samples. Read counts were then normalized with the trimmed mean of M-values (TMM) method^84^.

#### Exploration of covariates and model selection

We performed a PCA on the normalized read count matrix for each assay to identify high-variance components that explained at least 1% of the variance. Correlation tests were performed between the selected PCs and known covariates, and covariates with FDR<0.05 were used for the following steps. To select the final covariates, we first chose the covariates that are known to play a critical role for each assay as “a base model”: for instance, cell types, brain regions, and sex were selected as the base model for the Multi-omics cohort. We then applied an approach based on the Bayesian information criterion (BIC) to select the final covariates^43^. We examined the BIC changes in the linear regression model after adding a new covariate, which will be included if it can improve the mean BIC by at least 4.

#### Statistical test

With the selected covariates, the normalized read counts were modeled with the voomWithQualityWeights function from the limma package (v.3.38.3)^85^, which utilizes both sample-level and observational-level weights. We subsequently perform the test against the contrast between cell type or disease status using a linear mixed model to account for repeated measurements (i.e. 2 brain regions per individual) in the dream function^86^ of the variancePartition package^87^. Additionally, we determined the level of difference by estimating the proportion of true non-null tests Π_1_ ^88^ with the limma package.

Furthermore, we decomposed variation into known biological and technical factors with VarianceParition (v1.21.2)^87^. The analysis was performed by modeling the log_2_CPM with a linear mixed model and treating each variable as a random effect^86^. Results were summarized in terms of the fraction of total variation explained by each variable for each peak/transcripts.

### Identifying gene-enhancer links

For every gene, we considered all the enhancers within a 500 Kb window centered around the TSS (**Fig. 3c**) and fit a 10-fold cross-validation lasso model with the glmnet package (v 2.0.18)^89^. glmnet selects lambda to minimize the cross-validation prediction error and then selects lambda within 1 sd of the minimum. The enhancers with non-zero coefficients were selected as linked enhancers. For the genes that have only a single enhancer around the window, we performed a correlation test, and only the P_bonferroni_<0.01 were retained.

### Gene set enrichment analysis

To explore the function of a gene set, we collected functional gene sets from MSigDB 7.0^90^, human brain single-cell markers^38^, and synaptic gene ontology resource^91^. One-tailed Fisher exact tests were used to test the enrichment and significance.

To examine the genetic enrichment of gene sets, we used MAGMA (v 1.07b)^92^ with GWAS data^20,41,93–97^. Briefly, genes were padded by 35kb upstream and 10kb downstream, and the MHC region was removed due to its extensive linkage disequilibrium and complex haplotypes. The European panels from 1000 Genome Project phase 3 were used to estimate the Linkage disequilibrium (LD) ^98^.

### Partitioned heritability analysis

We partitioned heritability for DE peaks/TEns as well as top eSNPs to examine the enrichment of common variants in neuropsychiatric traits with stratified LD score regression (v.1.0.1)^22^ from a selection of GWAS studies^20,41,93–97^. Briefly, with the differential peaks/TEns as well as eSNPs, a binary annotation was created by marking all HapMap3 SNPs^99^ that fell within the peak or eSNPs and outside the MHC regions. LD scores were calculated for the overlapped SNPs using an LD window of 1cM using 1000 Genomes European Phase LD reference panel^98^. The enrichment was determined against the baseline model^22^. To enable comparisons of the regression coefficients across traits with a wide range of heritabilities, we chose to normalize it by the per-SNP heritability and named this adjusted metric the “heritability coefficient”. This is not the same as the “enrichment” also outputted by the software, since the heritability coefficient takes the aforementioned baseline into account and the “enrichment” does not.

### eQTL analysis

We used the MMQTL package to identify *cis*-QTLs for both genes and enhancers^51^. Briefly, Probabilistic Estimation of Expression Residuals (PEER) was used to determine a set of latent covariates to control for unknown biological and technical effects for the two brain regions, independently^100^. We use 30 PEER factors for ACC and 35 PEER factors for DLPFC based on the highest percentage of transcripts with eQTLs. The expression matrices were then adjusted for the selected covariates, the first 3 genetic principal components, and PEER factors. We performed a meta-analysis with the normalized expression matrices as well as the SNPs within 1Mb *cis*-window using a linear mixed model to maximize power^51^. P values for both GeQTL and EeQTL were corrected together for multiple testing using Storey and Tibshirani FDR correction^88^.

The proportion of null-hypotheses (Π_0_) was estimated by looking up the corresponding p values of the significant GTEx eQTL pairs (FDR<0.05) in our result with the qvalue package^88^. The non-null-hypothesis Π_1_=1-Π_0_ was subsequently determined. Additionally, we calculated the Spearman correlation coefficient between the effect sizes of our GeQTL and GTEx brain eQTL with all significant eQTLs (FDR<0.05). We performed a similar analysis to brain hQTL^54^ and caQTL^55^. Given that both analyses are done with hg19, we lifted over TEns to hg19 and found the overlapped peaks of the TEns positions. With the overlapped significant peak-SNP pairs, we calculated the Π_1_ using EeQTL p values for corresponding pairs.

To explore the functional enrichment of gene and enhancer eQTL, we performed enrichment analysis using Genomic Regulatory Elements and Gwas Overlap algoRithm (GREGOR)^56^ with our ChromHMM results. Briefly, we grouped genes and enhancers based on the differential expression status between neurons and non-neurons. The corresponding eSNPs were extended to all SNPs in high linkage disequilibrium (r^2^>0.7) and determined enrichment.

### MESC analysis

We used the mediated expression score regression (MESC) package^57^ to estimate SCZ heritability mediated by the *cis* genetic component of expression levels of gene, enhancer, or combined respectively. Based on the principle that expression mediated effect size introduces a linear relationship between the eQTL effect sizes and disease effect sizes, MESC determined the mediated effect through modeling GWAS summary statistics, LD scores, and eQTL effect sizes^57^. 1000 Genomes European Phase reference panel^98^ were used to compute LD scores. We retained only the SNPs that were from HapMap 3^99^ for this analysis. GCTA was used to estimate the *cis*-heritability (±500kb) of each transcript for two brain regions independently ^101^. Then we used the LASSO model from PLINK to estimate the eQTL size for each transcript. We performed a meta-analysis to combine the two brain regions to determine the expression score. Lastly, we estimated the SCZ heritability mediated by the expression of enhancer, gene, and combined transcripts.

### TWAS

The FUSION package was used for the gene-enhancer TWAS analysis^58^ for the two brain regions independently. For each brain region, only *cis*-heritable genes/enhancers (GCTA nominal p<0.05 and *cis*-heritability>0) were used for the following analysis. We built expression models using best *cis*-eQTL, Elastic-net regression, and LASSO regression with five-fold cross-validation^59^. The model with the best R^2^ was selected. Then we performed the association analysis with PGC3 SCZ GWAS summary statistics^41^. MHC regions were excluded for this analysis. The resulting p values of genes and enhancers are together Bonferroni corrected. Genes and enhancers with P_bonferroni_<0.05 were selected for downstream analysis.

We performed colocalization analysis using the bayesian estimator COLOC (v4.0.4)^64,65^ has been incorporated in the TWAS/FUSION pipeline. COLOC examine the posterior probability of five hypotheses: H0, no eQTL and no GWAS association; H1 and H2, associated to either eQTL or GWAS but not both; H3, eQTL and GWAS association but independent; H4, association. The analysis yields corresponding posterior probability PP0-PP4.

### FOCUS

To prioritize the potential casual transcripts for TWAS analysis, we performed statistical fine-mapping using FOCUS^66^. FOCUS models the correlation structure induced by LD and prediction weights in TWAS, and controls for certain pleiotropic effects. To account for the genes that are filtered by low *cis*-heritability, we collected gene prediction results from GTEx expression panels. For each gene, the model with the best accuracy was included. With gene/enhancer prediction results from two brain regions and the GTEx models, we performed FOCUS fine-mapping and got the 90% credible transcript sets.

## ACKNOWLEDGMENTS

We thank the computational resources and staff expertise provided by the Scientific Computing of the Icahn School of Medicine at Mount Sinai.

The CommonMind data sets were generated as part of the CommonMind Consortium supported by funding from Takeda Pharmaceuticals Company Limited, F. Hoffman-La Roche Ltd and NIH grants R01MH085542, R01MH093725, P50MH066392, P50MH080405, R01MH097276, RO1-MH-075916, P50M096891, P50MH084053S1, R37MH057881, AG02219, AG05138, MH06692, R01MH110921, R01MH109677, R01MH109897, U01MH103392, U01MH116442, project ZIC MH002903 and contract HHSN271201300031C through IRP NIMH. Brain tissue for the study was obtained from the following brain bank collections: The Mount Sinai/JJ Peters VA Medical Center NIH Brain and Tissue Repository, the University of Pennsylvania Alzheimer’s Disease Core Center, the University of Pittsburgh Brain Tissue Donation Program, and the NIMH Human Brain Collection Core. CMC Leadership: Panos Roussos, Joseph Buxbaum, Andrew Chess, Schahram Akbarian, Vahram Haroutunian (Icahn School of Medicine at Mount Sinai), Bernie Devlin, David Lewis (University of Pittsburgh), Raquel Gur (University of Pennsylvania), Chang-Gyu Hahn (Thomas Jefferson University), Enrico Domenici (University of Trento), Mette A. Peters, Solveig Sieberts (Sage Bionetworks), Stefano Marenco, Barbara K. Lipska, Francis J. McMahon (NIMH).

This work is supported by the National Institute on Aging, NIH grants R01-AG067025 (to P.R. and V.H.), R01-AG065582 (to P.R. and V.H.) and R01-AG050986 (to P.R.). Supported by the National Institute of Mental Health, NIH grants, R01-MH110921 (to P.R.), U01-MH116442 (to P.R. and V.H.), R01-MH125246 (to P.R.), R01-MH106056 (to P.R. and K.J.B.), R01-MH109897 (to P.R. and K.J.B.) and R01-MH121074 (to K.J.B.). Supported by the Veterans Affairs Merit grant BX002395 (to P.R.). P.D. was supported in part by NARSAD Young Investigator Grant 29683 from the Brain & Behavior Research Foundation. G.E.H. was supported in part by NARSAD Young Investigator Grant 26313 from the Brain & Behavior Research Foundation. J.B. was supported in part by NARSAD Young Investigator Grant 27209 from the Brain & Behavior Research Foundation.

## AUTHOR CONTRIBUTIONS

P.R. conceived of and designed the project. J.F.F. and P.R. designed experimental strategies for epigenome profiling of human postmortem tissue. J.F.F. prepared nuclei and performed FANS. J.F.F. and R.M. generated ATAC-seq data. P.A. generated the ChIP-seq and Multi-omics RNA-seq data. S.R. generated Hi-C data. S.R., J.M.V., M.B.F., K.G.T. and K.J.B. performed the CRISPR interference experiments. P.D. and P.R. designed analytical strategies. J.B. K.G. and P.D. conducted initial bioinformatics, sample processing and quality control for the Multi-omics cohort. G.E.H. and W.Z. conducted initial bioinformatics, sample processing and quality control for the CMC cohort. P.D. developed the computational scheme and performed the downstream analysis. B.Z. performed the MMQTL analysis. P.D. and P.R. wrote the manuscript with input from all authors.

## DECLARATION OF INTERESTS

The authors declare no competing interests.

## Data and code availability

The clinical information and raw data of the Multi-omics cohort including ATAC-Seq, RNA-Seq, H3K4me3/H3K27me3/H3K27ac ChIP-Seq and Hi-C are available via the AD Knowledge Portal (https://adknowledgeportal.org). The AD Knowledge Portal is a platform for accessing data, analyses, and tools generated by the Accelerating Medicines Partnership (AMP-AD) Target Discovery Program and other National Institute on Aging (NIA)-supported programs to enable open-science practices and accelerate translational learning. The data, analyses, and tools are shared early in the research cycle without a publication embargo on a secondary use. Data is available for general research use according to the following requirements for data access and data attribution (https://adknowledgeportal.org/DataAccess/Instructions). For access to content described in this manuscript see: http://doi.org/10.7303/syn25672193.

The processed data sets including ATAC-Seq/ChIP-Seq peaks, super-enhancer annotation, chromatin states, TEn annotation, differential analysis summary statistics, TEn-gene coordination, eQTL summary statistics, TWAS weights, and TEn identification pipeline are available on the Synapse platform at http://doi.org/10.7303/syn25716684. Further information and requests for reagents may be directed to the corresponding author/lead contact, Panos Roussos.

## Notes

http://doi.org/10.7303/syn25716684

### Competing Interest Statement

The authors have declared no competing interest.

