## Supplementary Notes for "Population-level variation of enhancer expression identifies novel disease mechanisms in the human brain"

### This Supplementary file includes:

Consortium author list

Supplementary Methods

References for Supplementary Methods

Extended Data Figs. 1-5

### CommonMind Consortium

The members of the CommonMind Consortium who are not listed in the primary author list are Andrew Chess^5,11^, Attila Gulyás-Kovács^5^, Bibi Kassim^4^, Eva Xia^4^, Joseph D Buxbaum^1,4^, Laura Sloofman^1,4^, Lizette Couto^4^, Mariana Amaro^1,4^, Marina Iskhakova^1,4^, Michael Breen^1,4^, Olivia Devillers^4^, Schahram Akbarian^4^, Shan Jiang^1,4^, Steven P Kleopoulos^1,4^, Yixian Ma^1,4^, Yungil Kim^1,4,5^, Sabina Berretta^12^, Ajeet Mandal^13^, Barbara K Lipska^13^, Francis McMahon^14^, Pavan K. Auluck^13^, Stefano Marenco^13^, Kelsey S Montgomery^15^, Mette A Peters^15^, Solveig K Sieberts^15^, Chang-Gyu Hahn^16^, Raquel Gur^17^, Jiebiao Wang^18^, Bernie Devlin^19^, David A Lewis^19^, Lambertus Klei^19^, Enrico Domenici^20^, Michele Filosi^20^, Roberto Visintainer^20^, Douglas M Ruderfer^21^, and Lide Han^21^.

^11^Institute for Genomics and Multiscale Biology, Icahn School of Medicine at Mount Sinai, New York, New York, USA

^12^Translational Neuroscience Laboratory, McLean Hospital, Belmont, MA, USA

^13^Human Brain Collection Core, ^14^Human Genetics Branch,

NIMH-IRP, Bethesda, MD, 20892, USA

^15^Sage Bionetworks, Seattle, Washington, USA

^16^Translational Neuropsychiatric Research Unit, Department of Psychiatry and Neuroscience, The Sidney Kimmel Medical College, Philadelphia, Pennsylvania, USA

^17^Neuropsychiatry Section, Department of Psychiatry, Perelman School of Medicine, University of Pennsylvania, Philadelphia, Pennsylvania, USA

^18^Department of Biostatistics, University of Pittsburgh, Pittsburgh, Pennsylvania, USA

^19^Department of Psychiatry, University of Pittsburgh School of Medicine, Pittsburgh, Pennsylvania, USA

^20^Laboratory of Neurogenomic Biomarkers, Centre for Integrative Biology (CIBIO), University of Trento, Trento, Italy

^21^Division of Genetic Medicine, Department of Medicine, Vanderbilt Genetics Institute, Vanderbilt University Medical Center, Nashville, TN, USA

### Supplementary Methods

#### CMC human postmortem brain samples

The CMC postmortem brain transcriptomic data consists of two non-overlapping cohorts. The CMC MSSM-Penn-Pitt Cohort (CMC MPP, used as discovery set), consists of samples of Anterior Cingulate Cortex (ACC) and Dorsolateral Prefrontal Cortex (DLPFC) from 545 postmortem brains accessed through the Mount Sinai NIH Brain Bank and Tissue Repository (MSSM), the University of Pennsylvania Brain Bank of Psychiatric illnesses and Alzheimer’s Disease Core Center (Penn), and the University of Pittsburgh NIH NeuroBioBank Brain and Tissue Repository (Pitt). The CMC HBCC Cohort (CMC_HBCC, used as a replicate set), consists of samples (ACC and DLPFC) from 229 postmortem brains accessed through the NIMH Human Brain Collection Core (HBCC). For both cohorts, dissections were performed at the corresponding brain banks and sent to Icahn School of Medicine at Mount Sinai for sample processing and library preparation. Postmortem tissue from schizophrenia (SCZ) cases was included if they met the diagnostic criteria in DSM-IV for SCZ or schizoaffective disorder after review of medical records, direct clinical assessments, and interviews with care providers. Individuals that had been diagnosed with, or had a history of other neuropsychiatric disorders (including bipolar disorder, and/or Parkinson’s disease), or acute neurological insults (anoxia, strokes, and/or traumatic brain injury), or Alzheimer’s disease were excluded from this analysis. Additionally, we excluded samples with age at death <17 and/or, PMI>48. All neuropsychological, diagnostic, and autopsy protocols were approved by the Mount Sinai and JJ Peters VA Medical Center Institutional Review Boards. The detailed subject information can be accessed through CMC Knowledge Portal (http://CommonMind.org).

#### FANS sorting of neuronal and non-neuronal nuclei

50 mg of frozen brain tissue was homogenized in chilled lysis buffer (0.32 M Sucrose, 5 mM CaCl_2_, 3 mM Magnesium acetate, 0.1 mM, EDTA, 10 mM Tris-HCl, pH 8, 1 mM DTT, 0.1% Triton X-100) and filtered through a 40 µm cell strainer. Filtered lysate was underlaid with sucrose solution (1.8 M Sucrose, 3 mM Mg(CH_3_COO)_2_, 1 mM DTT, 10 mM Tris-HCl, pH 8) and subjected to ultracentrifugation at 107,000 xg for 1 hour at 4°C. Pellets were resuspended in 500 µl DPBS containing 0.1% BSA. anti-NeuN antibody (1:1000, Alexa488 conjugated, Millipore, Cat# MAB377X) was added and samples incubated, in the dark, for 1 hr at 4°C. Prior to FANS sorting, DAPI (Thermoscientific) was added to a final concentration of 1 µg/ml. DAPI positive neuronal (NeuN+) and non-neuronal (NeuN-) nuclei were isolated using a FACSAria flow cytometer with FACSDiva Version 8.0.1 software (BD Biosciences).

#### Generation of RNA-seq libraries and sequencing

Multi-omics cohort. For RNA-seq, nuclei were sorted into 1.5 ml low-binding Eppendorf tubes containing Extraction buffer, a component of the PicoPure RNA Extraction kit (Arcturus, cat.# KIT0204). RNA was isolated following the manufacturer's instructions. This included an RNase-free DNase treatment step (Qiagen, cat.# 79254). Samples were eluted in RNase-free water and stored at -80°C until preparation of RNA-sequencing libraries using the SMARTer Stranded Total RNA-seq Pico Kit v1 (Takara Clontech Laboratories, cat.# 635005) according to the manufacturer’s instructions. Following the construction of the RNA-seq libraries, libraries were quantified by quantitative PCR (KAPA Biosystems, cat.# KK4873) and library fragment sizes estimated using High Sensitivity Tapestation D1000 ScreenTapes (Agilent, cat.# 5067-5584). RNA-seq libraries were subsequently sequenced on Hi-Seq2500 (Illumina) machines yielding 150 bp paired-end reads.

CMC MPP cohorts. Briefly, rRNA was depleted from about 1 µg of total RNA using Ribozero Magnetic Gold kit (Illumina/Epicenter, cat.# MRZG12324) to enrich polyadenylated coding RNA and non-coding RNA. The sequencing library was prepared using the TruSeq RNA Sample Preparation Kit v2 (Illumina, cat.# RS-122-2001). The insert size and DNA concentration of the sequencing library were determined on Agilent Bioanalyzer and Qubit, respectively. A pool of 10 barcoded libraries was layered on a random selection of two of the eight lanes of the Illumina flow cell at appropriate concentration and bridge amplified to ~250 million raw clusters. 100bp paired-end reads were obtained on a HiSeq 2500 (Illumina).

HBCC cohort. Briefly, rRNA was depleted from 1ug of RNA using the KAPA RiboErase protocol that is integrated into the KAPA Stranded RNA-seq Kit (KAPA Biosystems, cat.# KK8483). The insert size and DNA concentration of the sequencing library were determined on Fragment Analyzer Automated CE System (Advanced Analytical) and Quant-iT PicoGreen (ThermoFisher, cat.# P7589), respectively. A pool of 10 barcoded libraries was layered on a random selection of two of the eight lanes of the Illumina flow cell at appropriate concentration and bridge amplified to ~250 million raw clusters. 100bp paired-end reads were obtained by HiSeq 2500 (Illumina).

#### Generation of ATAC-seq libraries and sequencing

ATAC-seq libraries were generated using an established protocol[^1^](https://sciwheel.com/work/citation?ids=93501&pre=&suf=&sa=0). Briefly, 55,000 to 75,000 sorted nuclei were pelleted at 500 xg for 10 min at 4°C. Pellets were resuspended in transposase reaction mix (22.5 μL Nuclease Free H_2_O, 25 μL 2x TD Buffer; Illumina, cat.# FC-121-1030) and 2.5 μL Tn5 Transposase (Illumina, cat.# FC-121-1030) on ice and the reactions incubated at 37 °C for 30 min. Following incubation, samples were purified using the MinElute Reaction Cleanup kit (Qiagen cat.# 28204), and libraries generated using the Nextera index kit (Illumina cat. FC-121-1011). Following amplification, libraries were resolved on 2% agarose gels and fragments ranging in size from 100-1000 bp were excised and purified (Qiagen Minelute Gel Extraction Kit, Qiagen, cat.# 28604). Next, libraries were quantified by quantitative PCR (KAPA Biosystems, cat.# KK4873) and library fragment sizes estimated using Tapestation D5000 ScreenTapes (Agilent technologie, cat. 5067-5588). ATAC-seq libraries were subsequently sequenced on Hi-Seq2500 (Illumina) machines yielding 50 bp paired-end reads.

#### Generation of ChIP-seq libraries and sequencing

After the overnight incubation, the immunoprecipitation reactions were placed on a magnetic rack to remove unbound chromatin, washed twice with 200 µl of ChIP low salt wash buffer (20 mM Tris-HCl, pH 8.0, 0.1% SDS, 1% Triton X-100, 0.1% deoxycholate, 2 mM EDTA and 150 mM NaCl), twice with 200 µl ChIP high salt wash buffer (20 mM Tris-HCl (pH 8.0), 0.1% SDS, 1% Triton X-100, 0.1% deoxycholate, 2 mM EDTA and 500 mM NaCl), followed by elution in freshly prepared 30 µl of ChIP elution (100 mM sodium bicarbonate and 1% SDS) for 1.5 hours at 68°C on a ThermoShaker at 1000 rpm, RNAse A digestion for 15 minutes at 37˚C at 800 rpm. The immunoprecipitated DNA, along with the input controls, was purified using Phenol:Chloroform:Isoamyl Alcohol (25:24:1, v/v) (ThermoFisher Scientific, cat.# 15593-031), transferred to pre-spun phase lock tubes (Qiagen Maxtract, cat.# 129046) to obtain the aqueous layer. An overnight ethanol precipitation was performed by adding 10 μl of 3M sodium acetate/100 μl aqueous layer of ChIP DNA, 1 μl of LPA (linear polyacrylamide, Sigma #56575) and 1 μl Glycoblue (Invitrogen, cat.# AM9515) in 275 µl 100% ethanol.

NEBNext Ultra DNA Library Prep Kit (New England Biolabs, E7370) was used to construct NChIP libraries according to the manufacturer’s directions, followed by Pippin Size selection using 2% Agarose Gel cassettes (SAGE Science, cat.# HTC2010) and clean up with 1.8 volumes of SPRIselect beads (Beckman Coulter, cat.# B23317). All libraries were analyzed on an Agilent High Sensitivity D1000 TapeStation, and quantified using the KAPA Library Quantification Kit prior to sequencing. The constructed libraries, each containing a unique index, were pooled and sequenced using the Nova-seq platform (Illumina). Ultra-low input Native Chromatin Immunoprecipitation sequencing (ULI-NChIP-seq) libraries were prepared as follows. After isolating neuronal (NeuN+) and non-neuronal nuclei (NeuN- nuclei) by FANS, we performed ULI-NChIP assays, adapted from Brind’Amour, et al., 2015, which specifically does not require chromatin crosslinking, thereby increasing library complexity and reducing PCR artifacts. Briefly, nuclei were centrifuged at 500xg for 10 minutes at 4°C and re-suspended by gentle pipetting in residual smaller volumes of PBS/Sheath buffer. After DAPI counting of nuclei (Countess II, Life technologies), 100K-300K was distributed into Eppendorf tubes and 0.1% Triton-X-100/0.1% Na-Deoxycholate added. The chromatin was re-suspended and placed at room temperature for 5 minutes, followed by fragmentation with micrococcal nuclease (MNase, NEB, cat.# M0247S) for 5 min at 37°C on a ThermoShaker at 800 rpm to digest the chromatin to, predominantly, mononucleosomes. The MNase reaction was stopped by addition of 10% of the reaction volume of 100 mM EDTA (pipetted ~20x), followed by the addition of 1% Triton / 1% deoxycholate, (pipetted 5x) and placed on ice for at least 15 minutes. Samples were vortexed (medium setting) for ~ 30 seconds and complete NChIP buffer (20 mM Tris-HCl, pH 8.0, 2 mM EDTA, 15 mM NaCl, 0.1% Triton X-100, 1 EDTA-free protease inhibitor cocktail and 1 mM phenylmethanesulfonyl fluoride) was added to dilute the chromatin to <25% of the immunoprecipitation reaction volume, and rotated at 4°C for 1 hour. After the incubation, chromatin was vortexed (medium setting) for ~30 seconds and 5% input controls removed for DNA extraction.

To avoid non-specific binding, chromatin was next precleared by adding 10 µl/ reaction of a pre-washed 1:1 ratio of protein A to protein G Dynabeads (Life Technologies), rotating the chromatin-protein A/ protein G magnetic bead mixture for 3 hours at 4°C. Antibody-bead complexes were prepared as follows: ChIP-grade histone H3K27ac antibodies (Active Motif, cat.# 39133, pAB), H3K4me3 (Cell Signaling, cat.# 9751), or H3K27me3 (Cell Signaling, cat.# 9733) were added to prewashed protein A/protein G Dynabeads resuspended in NChIP buffer, and the antibody-bead complexes formed by rotating the antibody and beads for 2 hours at 4°C. After the incubations, precleared chromatin and antibody-bead complexes were placed on a magnetic rack and the precleared chromatin transferred to new Eppendorf tubes, while the antibody-bead complexes were resuspended in sufficient volume of NChIP buffer to add 10 µl per MNase reaction. The chromatin was immunoprecipitated with the antibody-bead complexes at 4°C overnight while rotating.

#### Generation of Hi-C libraries and sequencing

Hi-C data was generated from frozen postmortem human brain tissue using the in situ Hi-C protocol[^2^](https://sciwheel.com/work/citation?ids=48490&pre=&suf=&sa=0) with the following modifications. Frozen brain tissue was thawed at room temperature (RT) and dounce homogenized in HBSS (Hank’s balanced salt solution). Homogenized tissue was fixed with 0.5% formaldehyde for 10 min and then quenched with 0.125 M glycine for 5 min at RT. Cross-linked tissue was then placed on ice for a further 15 min to quench crosslinking completely. Samples were centrifuged at 800 xg for 10 min at 4°C and pellets resuspended in lysis buffer (0.32 M Sucrose, 5 mM CaCl_2_, 3 mM Mg(CH_3_COO)_2_, 0.1 mM EDTA, 10 mM Tris-HCl, pH 8, 1 mM DTT, 0.1 % Triton X-100, 1x Roche cOmplete mini EDTA-free protease inhibitor tablet (Roche cat.# 4693159001) to isolate cross-linked nuclei. Neuronal nuclei were FANS sorted using the Alexa488 conjugated anti-NeuN antibody (1:1000) (Millipore, cat.# MAB377X) using a BD FACS Aria II sorter. FANS sorted neuronal (NeuN+) and non-neuronal (NeuN-) nuclei were pelleted at 2500 xg for 5 min at 4°C, frozen on dry ice for 20 min, and then stored at -80°C.

Approximately 1 M crosslinked neuronal and non-neuronal nuclei were thawed on ice, washed with ice-cold 1x CutSmart buffer (New England Biolabs (NEB), cat. B7204S) and split into 4 aliquots to generate technical replicate libraries per sample, with 250k nuclei per library. Nuclei were pelleted at 2500 xg for 5 min at 4°C, and resuspended in 342 µl 1x CutSmart buffer and then conditioned with 0.1% SDS at 65°C for 10 min. Nuclei were immediately placed on ice and the SDS quenched with 1% Triton X-100. Chromatin was digested with 100 U of the 4 base pair cutter MboI (NEB, cat.# R0147L) overnight at 37°C with shaking at 400 rpm. MboI was heat-inactivated at 65°C for 20 min, and then nuclei were cooled down on ice. MboI cut sites were end-labeled with biotin by adding 52 µl of biotin fill-in reaction mix (15 µl of 1 mM biotin-14-dATP (Jena Bioscience, cat.# NU-835-BIO14-L), 1.5 µl each of 10 mM dCTP, dGTP, dTTP (Sigma-Aldrich, cat.# DNTP10-1KT), 10 µl of 5 U/µl Klenow DNA Pol I (NEB, cat.# M0210L), 22.5 µl of 1x CutSmart buffer) and shaking at 37°C for 1.5 h at 400 rpm. Blunt ended sites were proximity ligated by adding 948 µl ligation reaction mix (150 µl of 10x T4 DNA ligase buffer (NEB, cat.# B0202S), 125 µl of 10% Triton X-100, 15 µl of 10 mg/ml BSA, 10 µl of 400 U/µl T4 DNA ligase (NEB, cat.# M0202L), 648 µl ddH_2_0) and rotating tubes, end-over-end, at RT for 4 h. Nuclei were reverse crosslinked with 100 µl proteinase K (10 mg/ml) overnight at 65°C.

Proximity ligated DNA was purified through phenol:chloroform extraction and sodium acetate/ethanol precipitation. Purified DNA was sheared using a Covaris S220 sonicator to generate a peak size of 400 bp with the following settings (peak incident power: 140 W, duty cycle: 10%, cycles per burst: 200, time: 55 sec). Biotin-labeled ligation junctions were purified with Dynabeads MyOne Streptavidin C1 beads (ThermoFisher, cat.# 65001) by incubating for 1 hr at RT. Illumina compatible libraries were prepared from the sonicated and streptavidin bead immobilized DNA using the NEBNext Ultra II Library prep kit (NEB, cat.# E7645L), following manufacturer’s instructions, by amplifying libraries for 6-10 PCR cycles. Libraries were purified by 2-sided size selection (300-800 bp) using Ampure XP beads (Beckman Coulter, cat.# A63881). All libraries were analyzed on a TapeStation using Agilent D5000 ScreenTapes (Agilent technologies, cat.# 5067-5588), and quantified using the KAPA Library Quantification Kit (KAPA Biosystems, cat.# KK4873) prior to sequencing. Uniquely barcoded Hi-C libraries were pooled and deep sequenced on the Illumina NovaSeq S4 platform (Illumina) obtaining 100 bp paired-end reads.

#### Whole-genome Sequencing data processing

Whole-genome sequencing (WGS) data were obtained from AMP-AD MSBB cohort[^3^](https://sciwheel.com/work/citation?ids=5733392&pre=&suf=&sa=0). The paired-end 150 bp reads were aligned to the human hg38 reference genome with Burrows–Wheeler Aligner (bwa) using the bwa-mem algorithm[^4^](https://sciwheel.com/work/citation?ids=10532255&pre=&suf=&sa=0). Duplicates were marked and discarded by Picard. Then the bam files were realigned around indels, and base recalibrated with GATK. For each individual, variants were called using GATK HaplotypeCaller. The individual files were then jointly genotyped using GATK GenotypeGVCFs and generated a multi-sample VCF. Finally, quality metrics for each variant were determined through Variant Quality Score Recalibration (VQSR).

#### SNP array data processing

For the CMC-MPP cohort, genotyping was performed on the Illumina Infinium HumanOmniExpressExome 8 v 1.1b chip (Catalog #: WG-351-2301) using the manufacturer’s protocol. For the CMC HBCC cohort, genotyping was performed on one of 3 different Illumina gene chips: HumanHap650Y, Human1M-Duo, and HumanOmni5M-Quad, according to the manufacturer’s instructions.

Genotyping QC and imputation proceeded separately by gene chipset as previously described[^5^](https://sciwheel.com/work/citation?ids=8218833&pre=&suf=&sa=0). Briefly, markers with: zero alternate alleles, genotyping call rate ≤ 0.98, Hardy-Weinberg P value < 5x10^-5^, and individuals with genotyping call rate < 0.90 were removed by PLINK[^6^](https://sciwheel.com/work/citation?ids=431749&pre=&suf=&sa=0). Then, samples were imputed to HRC (r1.1 2016)[^7^](https://sciwheel.com/work/citation?ids=2311631&pre=&suf=&sa=0). Strands, alleles, position, reference/alternate allele assignments and frequencies were checked; and SNPs that with: A/T & G/C alleles and minor allele frequency (MAF) > 0.4, differing alleles, > 0.2 allele frequency difference between the genotyped samples and the HRC samples, and not in reference panel were removed. Imputation was performed with the Michigan Imputation Server[^8^](https://sciwheel.com/work/citation?ids=2094306&pre=&suf=&sa=0) using Eagle (v2.3)[^9^](https://sciwheel.com/work/citation?ids=3224851&pre=&suf=&sa=0). All imputation was performed using the GRCh37, the genotype data were subsequently lifted over to GRCh38 for downstream integration with functional genomics assays.

#### RNA-seq data processing

Multi-omics cohort. The trimmed reads were aligned to human genome hg38 (GRCh38) using STAR (2.7.2a) aligner[^10^](https://sciwheel.com/work/citation?ids=49324&pre=&suf=&sa=0), where the allelic alignment bias was corrected by WASP[^11^](https://sciwheel.com/work/citation?ids=814508&pre=&suf=&sa=0). Gene expression was quantified using RSEM (v1.3.1) tools[^12^](https://sciwheel.com/work/citation?ids=707264&pre=&suf=&sa=0) with GENCODE V30 as a reference and summarized at the gene level. Duplication and GC content levels were estimated by Picard tools (v2.2.4). Quality control (QC) metrics were collected by RNA-seq QC (v1.1.8)[^13^](https://sciwheel.com/work/citation?ids=463338&pre=&suf=&sa=0). The RNA-seq libraries were deeply sequenced and have acceptable values for total reads (mean 1.82x10^8^, sd ±9.5x10^7^), final read pairs (mean 1.48x10^8^ sd ±7.44x10^7^), intergenic rate (mean 11.7%, sd ±1.86%), intronic rate (mean 59.3%, sd ±3.38%), and ribosomal RNA rate (mean 0.0906%, sd ±0.0631%). We checked the sex compatibility using the expression of two sex-specific genes XIST and RPS4Y1.

RNA-Seq cell type deconvolution analysis. We used dTangle (v2.0.9)[^14^](https://sciwheel.com/work/citation?ids=6017166&pre=&suf=&sa=0), a method built on the linear mixing model of linear-scale expressions of known marker genes, to estimate the cell type composition of RNA-seq data. Human brain single-cell markers including five brain cell types: glutamatergic neurons, GABAergic neurons, astrocytes, oligodendrocytes, and microglia were used as the reference[^15^](https://sciwheel.com/work/citation?ids=4709763&pre=&suf=&sa=0). As expected, the neuronal cells are enriched for GABAergic and glutamatergic, while the non-neuronal cells are strongly enriched for oligodendrocytes and followed by astrocytes and microglia (**Extended Data Fig. 1a**).

CMC Cohort. RNA-seq data were processed using the Multi-omics RNA-seq pipeline, as described above. The RNA-seq libraries were deeply sequenced and have acceptable values for total read (mean 5.18x10^7^, sd ±2.61x10^7^), final read pairs (mean 4.78x10^7^, sd ±2.35x10^7^), intergenic rate (mean 5.75%, sd ±2.69%), intronic rate (mean 38.6%, sd ±10.3%), ribosomal RNA rate (mean 0.103%, sd ±0.299%), and RIN (mean 7.41, sd ±0.977 ).

#### ATAC-seq data processing

Alignment. The trimmed reads were aligned to human genome hg38 (GRCh38) using STAR (2.7.0e) aligner[^10^](https://sciwheel.com/work/citation?ids=49324&pre=&suf=&sa=0), with customized parameters. We restricted the intron length (--alignIntronMax 1), number of mismatches per pair ( --outFilterMismatchNmax 100), alignment score ( --outFilterScoreMinOverLread 0.66) and number of matched bases ( --outFilterMatchNminOverLread 0.66) in local alignment mode (-alignEndsType Local), and controlled allelic alignment bias with WASP[^11^](https://sciwheel.com/work/citation?ids=814508&pre=&suf=&sa=0). Reads mapped to multiple loci detected by samtools (v0.1.19), duplicated reads marked by Picard (v2.2.4), and mitochondria alignments were filtered.

Peak calling. We called peaks for each cell type independently. For each cell type, bam files from each sample were subsampled to the same sequencing depth and then merged. We used model-based Analysis of ChIP-seq (MACS, v2.1)[^16^](https://sciwheel.com/work/citation?ids=57981&pre=&suf=&sa=0) to call peaks with a smoothing window of 200bps ( --shift -100 --extsize 200 --nomodel) and FDR of 0.01 (-q 0.01). Peaks overlapped with ENCODE blacklist regions[^17^](https://sciwheel.com/work/citation?ids=7132831&pre=&suf=&sa=0) were discarded, yielding 222,746 neuronal and 136,171 non-neuronal OCRs. To confirm our results, we determined the Jaccard index between our peaks and previously published results[^18^](https://sciwheel.com/work/citation?ids=5486493&pre=&suf=&sa=0), which is the intersection of base pairs divided by the union of base pairs (**Extended Data Fig. 1b**).

Quantification. A consensus peak set of both cell types was generated for quantification and differential analysis. We counted the number of reads that overlapped with the consensus peaks using featureCounts function in RSubread (v 1.6.3)[^19^](https://sciwheel.com/work/citation?ids=148598&pre=&suf=&sa=0).

QC. We collected the following information for each sample for QC. 1) total number of initial reads; 2) the number of uniquely mapped reads; 3) the fraction of reads that were uniquely mapped; 4) further mappability-related metrics from the STAR aligner; 5) GC content, 6) insert and duplication metrics from Picard; 6) the rate of reads mapping to the mitochondrial genome; 7) the PCR bottleneck coefficient (PBC), which approximates library complexity as uniquely mapped non-redundant reads, divided by the number uniquely mapped reads; 8) the relative strand cross-correlation coefficient (RSC) and the normalized strand cross-correlation coefficient (NSC), which are metrics that use cross-correlation of stranded read density profiles to evaluate the sample quality independently of peak calling; and, 10) finally, the fraction of reads in peaks (FRiP), which is the fraction of reads that fall in called peaks within backlist regions or not. The ATAC-seq libraries have acceptable values for: total reads (mean 5.61x10^7^, sd ± 9.2x10^6^), final read pairs (mean 3.63x10^7^, sd ± 8.07x10^6^), mitochondrial alignment rate (mean 1.59%, sd ± 0.61%), fraction of reads in non-blacklist peaks (mean 17.6%, sd ± 4.12%), and mean GC content (mean 51.4%, sd ± 3.58%).

#### ChIP-seq data processing

H3K4me3, H3K27ac, H3K27me3 ChIP-seq, and corresponding input files were aligned to human genome hg38, similar to our ATAC-seq pipeline, above. The resulting bam files were subsampled to the same sequencing depth and merged for each cell type. We called peaks using MACS with a smoothing window of 150bps ( --shift -75 --extsize 150 --nomodel), FDR of 0.01 (-q 0.01), and input as control. For H3K27me3, we called broad peaks (--broad). Peaks overlapped with ENCODE blacklist regions[^17^](https://sciwheel.com/work/citation?ids=7132831&pre=&suf=&sa=0) were discarded, yielding 76,905 H3K4me3, 312,658 H3K27ac, and 83,865 H3K27me3 neuronal peaks, as well as 96,015 H3K4me3, 227,155 H3K27ac, and 68,457 H3K27me3 non-neuronal peaks. Comparing H3K4me3 and H3K27ac peaks to reported peaks[^20^](https://sciwheel.com/work/citation?ids=5579318&pre=&suf=&sa=0) by determining the Jaccard index (**Extended Data Fig. 1b**), we observed high concordance between cell types. For H3K27me3 peaks, we compared our cell-type-specific peaks with brain DLPFC bulk H3K27me3 peaks[^21^](https://sciwheel.com/work/citation?ids=48808&pre=&suf=&sa=0). A consensus peak set was generated for quantification and differential analysis. We quantified the number of reads overlapped with the consensus peaks using featureCounts function in RSubread (v 1.6.3)[^19^](https://sciwheel.com/work/citation?ids=148598&pre=&suf=&sa=0). The QC metrics were collected as for the ATAC-seq data, above.The H3K4me3 ChIP-seq libraries have acceptable values for: total read (mean 1.18x10^8^, sd ± 3.6x10^7^), final read pairs (mean 5.98x10^7^, sd ± 1.87x10^7^), fraction of reads in non-blacklist peaks (mean 83.1%, sd ± 9.23%), NSC (mean 2.14, sd ± 0.24), and RSC (mean 1.46, sd ± 0.135). The H3K27ac ChIP-seq libraries have acceptable values for: total read (mean 1.04x10^8^, sd ± 2.45x10^7^), final read pairs (mean 7.23x10^7^, sd ± 1.91x10^7^), fraction of reads in non-blacklist peaks (mean 71.5%, sd ± 5.76%), NSC (mean 1.19, sd ± 0.0425), and RSC (mean 3.25, sd ± 0.748). The H3K27me3 ChIP-seq libraries have acceptable values for: total read (mean 1.78x10^8^, sd ± 8.03x10^7^), final read pairs (mean 1.35x10^8^, sd ± 5.04x10^7^), fraction of reads in non-blacklist peaks (mean 44.2%, sd ± 1.07%), NSC (mean 1.09, sd ± 0.0377), and RSC (mean 3.09, sd ± 1.09).

#### Super enhancer identification

We identified super-enhancers with H3K27ac ChIP-seq peaks and bam files using the ROSE pipeline[^22^](https://sciwheel.com/work/citation?ids=48468&pre=&suf=&sa=0). Briefly, enhancer peaks were stitched together if they are located within 12.5 kb of each other and don’t have multiple active promoters in between. The stitched peaks were then ranked according to increasing H3K27ac signal intensity. We identified 2,049 neuronal and 1,946 non-neuronal super-enhancers, respectively.

#### Chromatin states annotation

We implemented a multivariate Hidden Markov Model model (ChromHMM)[^23^](https://sciwheel.com/work/citation?ids=590356&pre=&suf=&sa=0) to systematically annotate the combinational effect of different histone modifications. The ChromHMM model is trained by virtually concatenating histone marks H3K4me3, H3K27ac, and H3K27me3 in both cell types that merged across all individuals that subsampled to uniform depths. Reads were shifted from 5’ to 3’ direction by 100 bp for all the samples. Read counts were then computed in 200bp non-overlapping bins across the genome. Each bin was binarized into 1 or 0 by the Poisson model with a p-value threshold of 10^-4^. We have trained the model with merged data using six states which captured all the key interactions from our data. Lastly, we obtained the chromatin states with the trained model and corresponding binarized files as input (**Extended Data Fig. 2a**).

Our models agreed well with the published brain DLPFC region 11 chromatin states model [^21^](https://sciwheel.com/work/citation?ids=48808&pre=&suf=&sa=0) (**Extended Data Fig. 2b**).

#### Hi-C data analysis

Hi-C data were aligned using the HiC-Pro strategy[^24^](https://sciwheel.com/work/citation?ids=1036103&pre=&suf=&sa=0). Briefly, paired-end reads were mapped independently to the human genome hg38 using bowtie2 in stringent mode with parameters (‘--very-sensitive -L 20 --score-min L,-0.6,-0.2 --end-to-end’)[^25^](https://sciwheel.com/work/citation?ids=48791&pre=&suf=&sa=0). Then, the chimeric reads that failed to align were trimmed after ligation sites (MboI ‘GATCGATC’) and mapped to the genome. All the aligned reads from both ends were then merged based on read names and mapped to MboI restriction fragments using hiclib package[^26^](https://sciwheel.com/work/citation?ids=48630&pre=&suf=&sa=0). Next, self-circles, dangling ends, PCR duplicates, and genome assembly errors were discarded. Samples of the same cell type were merged. We binned the interaction matrix at different resolutions and corrected it with iterative correction (ICE) for downstream analysis.

Chromatin loops were called with HICCUPS[^27^](https://sciwheel.com/work/citation?ids=4967221&pre=&suf=&sa=0) for the two different cell types independently. First, we converted the filtered interaction files into juicer format with juicertools. Chromatin loops were called using juicer HICCUPS with bin sizes iterated from 10kb to 25kb by 1kb intervals and parameters ‘-k VC_SQRT -p 1 -i 3’. Only reproducible loops were retained and the highest resolution of the overlapping loops was used.

Topological associated domains (TADs) were identified with Topdom[^28^](https://sciwheel.com/work/citation?ids=1308064&pre=&suf=&sa=0) at 10K resolution and a 200Kb window size.

#### Genetic variants concordance analysis

To check for sample mix-ups, we evaluated the genetic variants between WGS genotyping and RNA-seq, ATAC-seq, and ChIP-seq samples. For RNA-seq, ChIP-seq, and ATAC-seq, we called genotypes with GATK (v3.5.0)[^29^](https://sciwheel.com/work/citation?ids=56163&pre=&suf=&sa=0). We performed i) indel-realignment, ii) base score recalibration and iii) joint genotype calling across all samples. Variants with a phred-scaled confidence threshold <10, within ENCODE blacklisted regions of the genome[^17^](https://sciwheel.com/work/citation?ids=7132831&pre=&suf=&sa=0), outside of dbSNP v151, minor allele frequencies (MAF) <25%, and clustered variants were not used. The genotype concordance amongst samples was quantified using both the fraction of concordant genotype calls and the kinship coefficient from KING (v1.9)[^30^](https://sciwheel.com/work/citation?ids=1433823&pre=&suf=&sa=0), where the two methods give consistent results.

#### CAGE-seq data processing

CAGE-seq data of four brain lobes including occipital, frontal, temporal, cerebellum were downloaded from the published result (NCBI Bioproject PRJNA273171)[^31^](https://sciwheel.com/work/citation?ids=2297640&pre=&suf=&sa=0). We mapped the clean reads to the human reference genome hg38 with STAR (2.5.3a) aligner. The BAM files of the four brain regions were merged for downstream analysis.

### Extended Data Figures

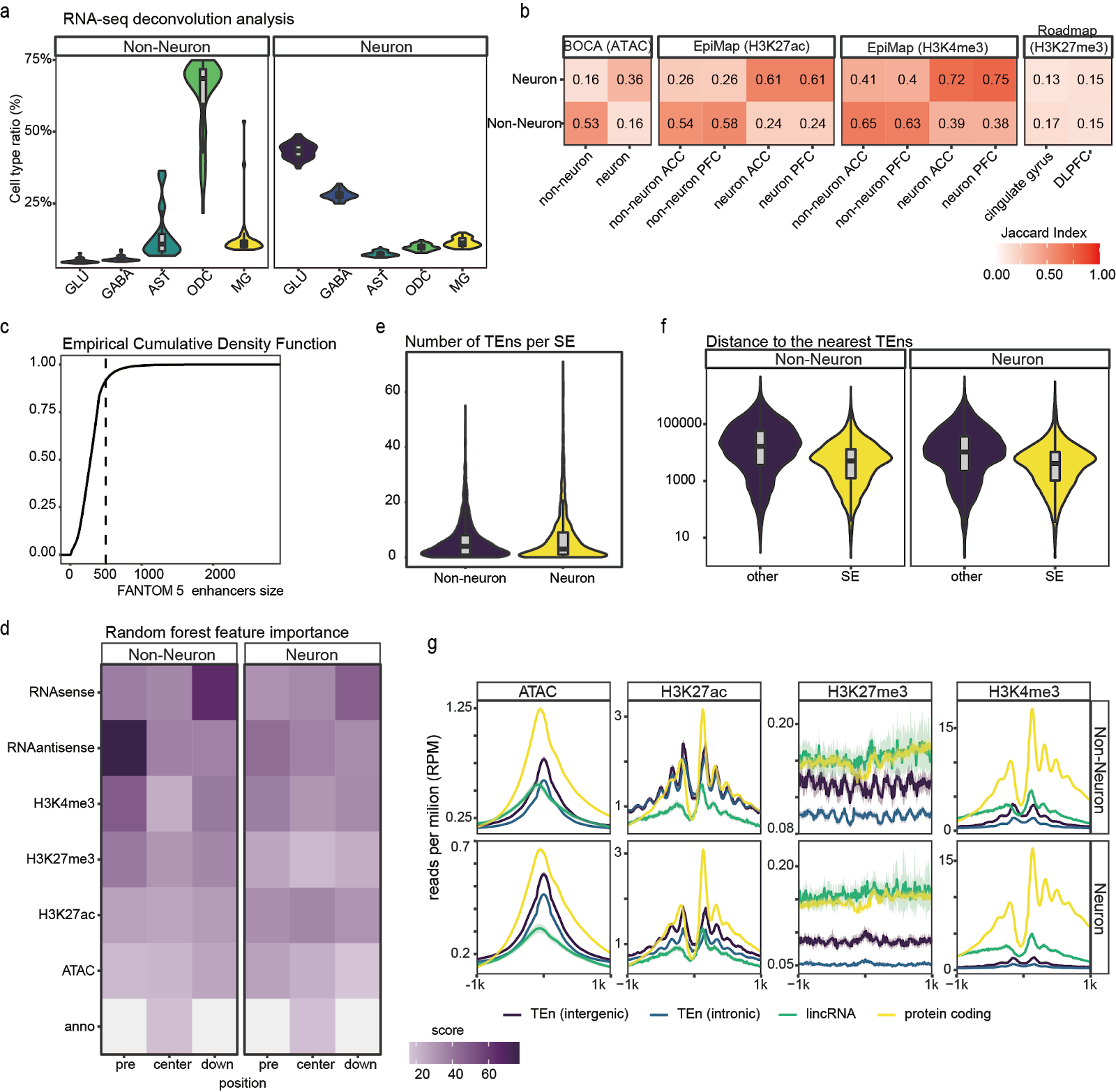

###### **Extended Data Fig. 1 | TEns identification**. **a**, The distribution of deconvolved RNA-seq cell type distribution for each sample. **b**, Jaccard index between the Multi-omics peaks and previous reports. cell-type-specific ATAC-seq/H3K4me3/H3K27ac peaks were compared to our previous reports of corresponding assays[^18,20^](https://sciwheel.com/work/citation?ids=5486493,5579318&pre=&pre=&suf=&suf=&sa=0,0). H3K27me3 was compared to Roadmap H3K27me3 peaks[^21^](https://sciwheel.com/work/citation?ids=48808&pre=&suf=&sa=0). PFC indicates the prefrontal cortex. **c**, Empirical density distribution of FANTOM5 enhancer size; based on the curve, we chose 500bp as the TEn size. **d**, Feature importance heatmap from random forest models. **e**, Distribution of TEn numbers per super-enhancer in the two cell types. **f**, Distance to the nearest TEns for every TEn within or outside of super-enhancer regions (p<10^-16^ for both cell types, two-sided Wilcoxon test). **g**, Epigenomic profiles of expressed TEns, and promoters of protein-coding genes and lincRNAs. Shadow indicates a 95% confidence interval.

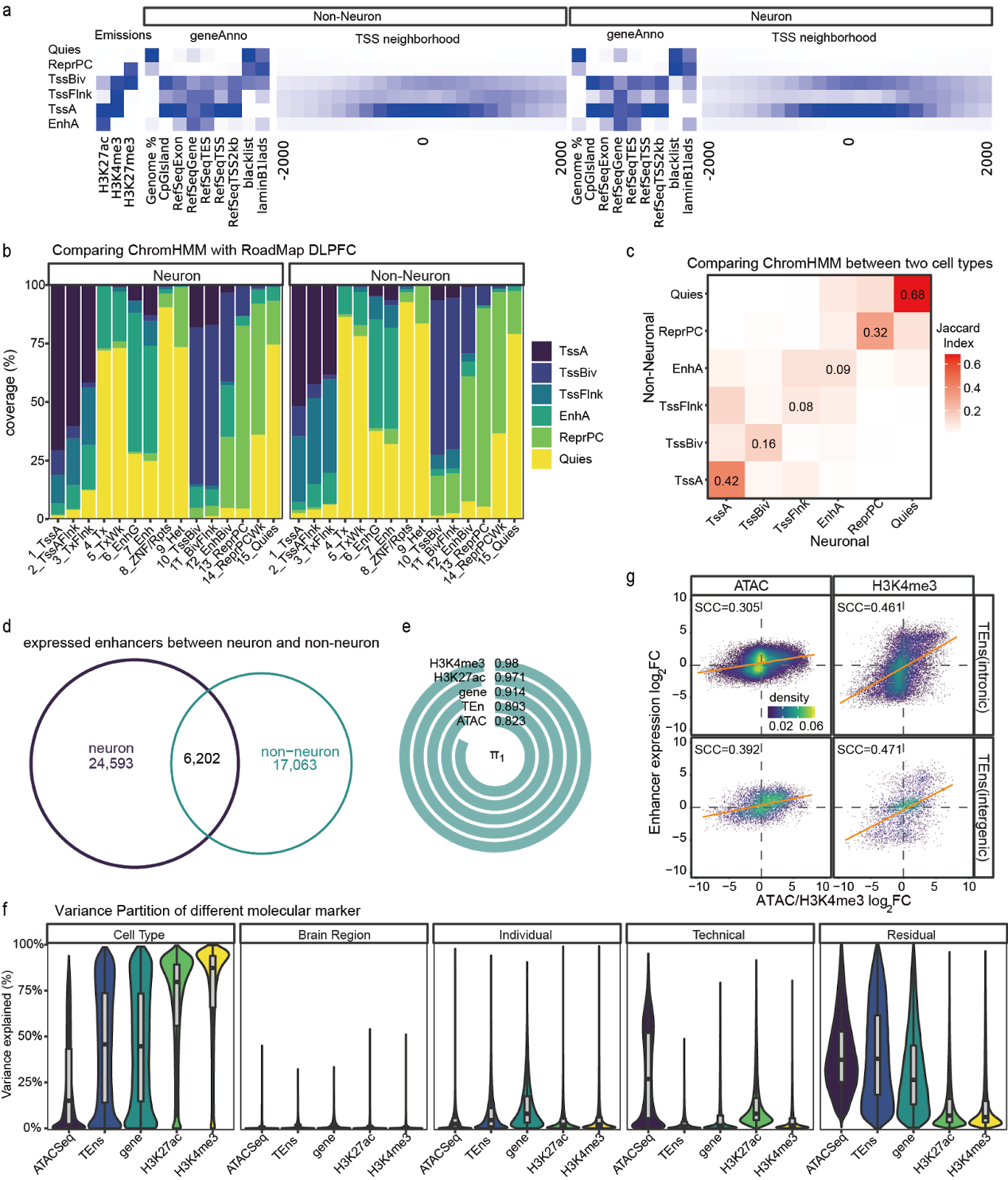

###### **Extended Data Fig. 2 | Differential expression/activity between neuronal and non-neuronal cells. a**, ChromHMM identified 6 chromatin states including EnhA (active enhancer), TssA (active promoters), TssBiv (bivalent promoters), TssFlnk (promoter flanking region), ReprPC (polycomb repression region), and Quies (other regions). **b**, Comparing the cell-type-specific Chromatin states with the roadmap DLPFC result. **c**, Jaccard index between neuronal and non-neuronal chromatin states. Compared to active promoters (TssA), and polycomb repressed regions (ReprPC), active enhancers (EnhA) are remarkably more different between cell types. **d**, Overlap of expressed TEns between neuronal and non-neuronal cells. **e**, π_1_ statistics of differential activity between the two cell types of different molecular markers. **f**, Violin plot shows the variance explained by different factors for the five markers. **g**, cell-type-specific effect size (log_2_ fold change) between TEns and physically overlapping ATAC-seq/H3K4me3 peaks are highly consistent for both intergenic (N_ATAC_=8,309, N_H3K4me3_=4,910) and intronic (N_ATAC_=46,509, N_H3K4me3_=22,496) TEns. SCC represents the Spearman correlation coefficient (all p<10^-16^).

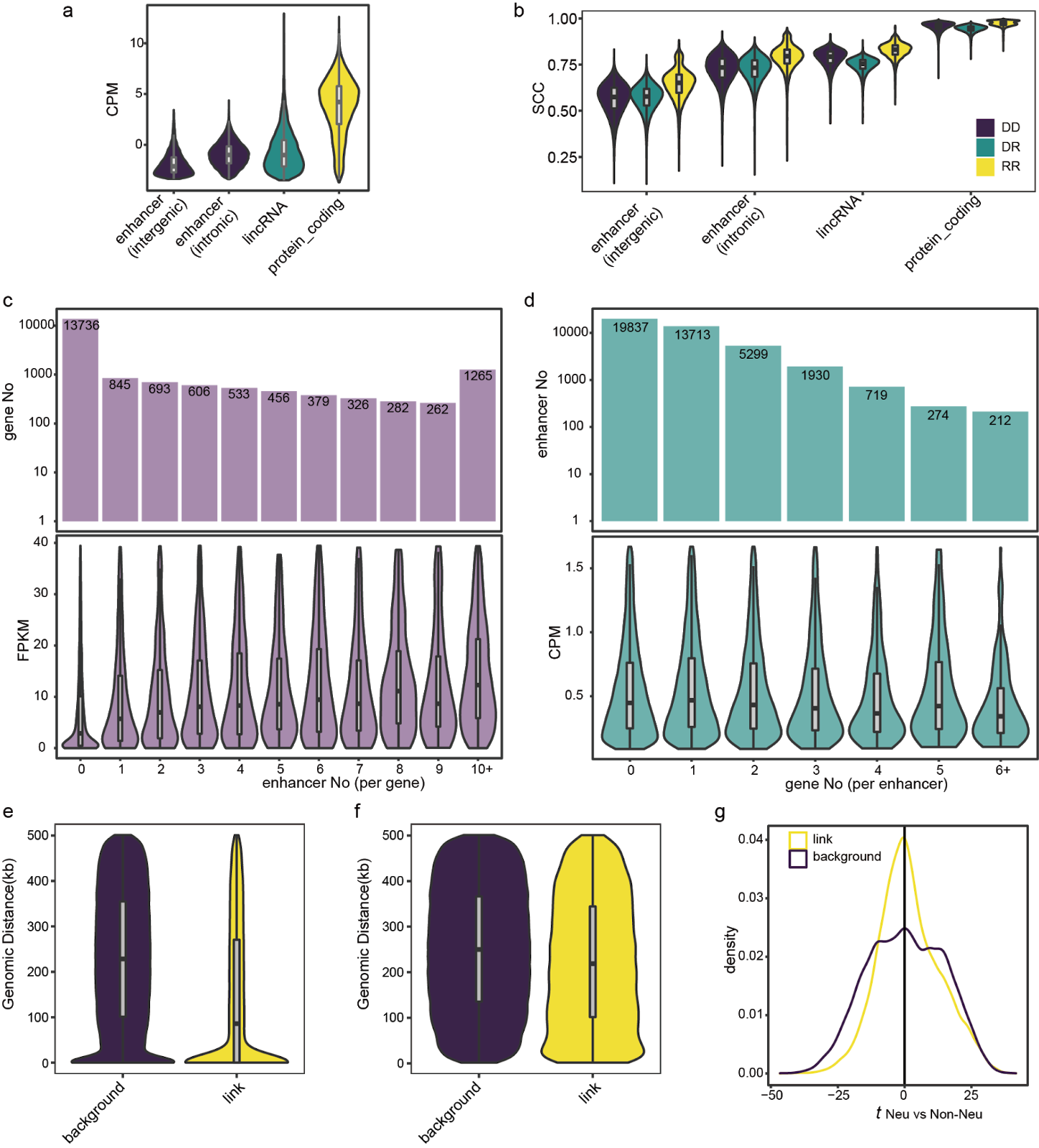

###### **Extended Data Fig. 3 | enhancer-gene expression coordination. a**, expression level of different classes of genes and enhancers. **b**, Pairwise Spearman correlation coefficient (SCC) between samples of different classes of genes/enhancers of discovery vs discovery (DD), discovery vs replicate (DR), and replicate vs replicate (RR). **c**, Distribution of gene FPKM and counts for linked enhancers. Linked genes have a significantly higher gene expression (FPKM, two-sided Wilcoxon test, p<10^-16^). **d**, Distribution of enhancer CPM and for counts linked genes. Linked enhancers do not have a higher expression (CPM, two-sided Wilcoxon test, p=0.96). **e**, Distance between enhancers to genes of linked group and background (N_link_=35,964, N_background_=241,040). **f**, Distance between enhancers to genes of linked group and background, physically-overlapped gene-enhancer pairs were excluded (N_link_=22,920, N_background_=220,062). **g**, Neuronal vs non-neuronal t statistics for linked and not linked enhancers. Linked enhancer have a significantly higher value (KS test, p<10^-16^)

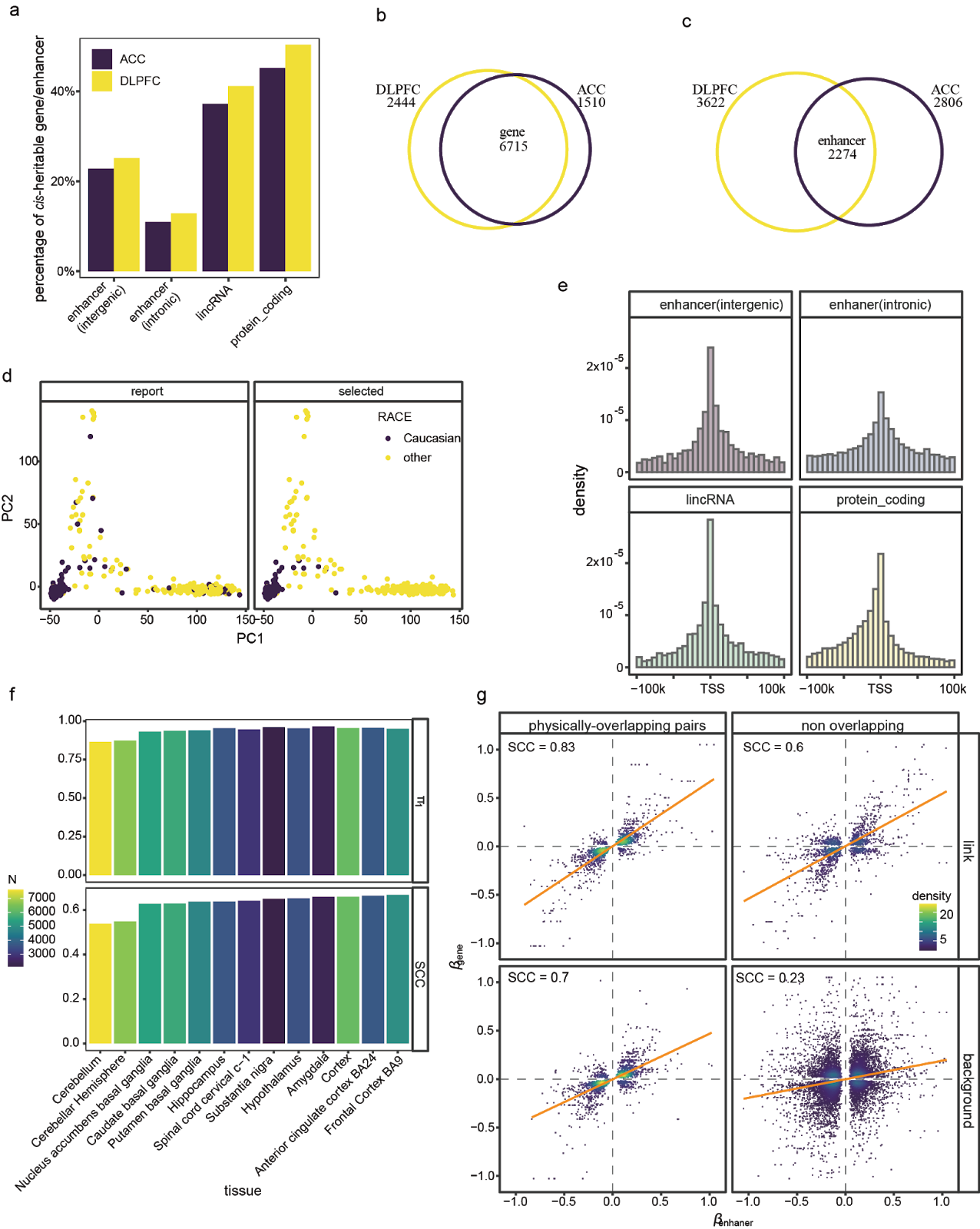

###### **Extended Data Fig. 4 | gene and enhancer eQTL. a**, Percentage of genes/enhancers that are cis-heritable (p<0.05 and cis-heritability>0). **b**, Overlap of cis-heritable genes between ACC and DLPFC. **c**, Overlap of cis-heritable enhancers between ACC and DLPFC. **d**, Dot plots illustrate the first two genetic ancestry principal components (PCs) for individuals reported (report) to have European ancestry and the individuals selected (select) to have European ancestry based on sd to the center. **e**, The distribution of genomic distances from eSNPs to the TSSs for different classes of transcripts. **f**, The replication of reported eQTL in our analysis. π_1_ values (proportion of true positive p values) for reported significant GTEx eQTL in our gene eQTLs. SCC values are the Spearman correlation coefficients of significant eQTL effect sizes between GTEx eQTL and corresponding pairs in our data. The bar color represents the number of unique genes used. **g**, The allelic genetic effect between gene and target enhancers considering the gene-enhancer physically-overlapping effect.

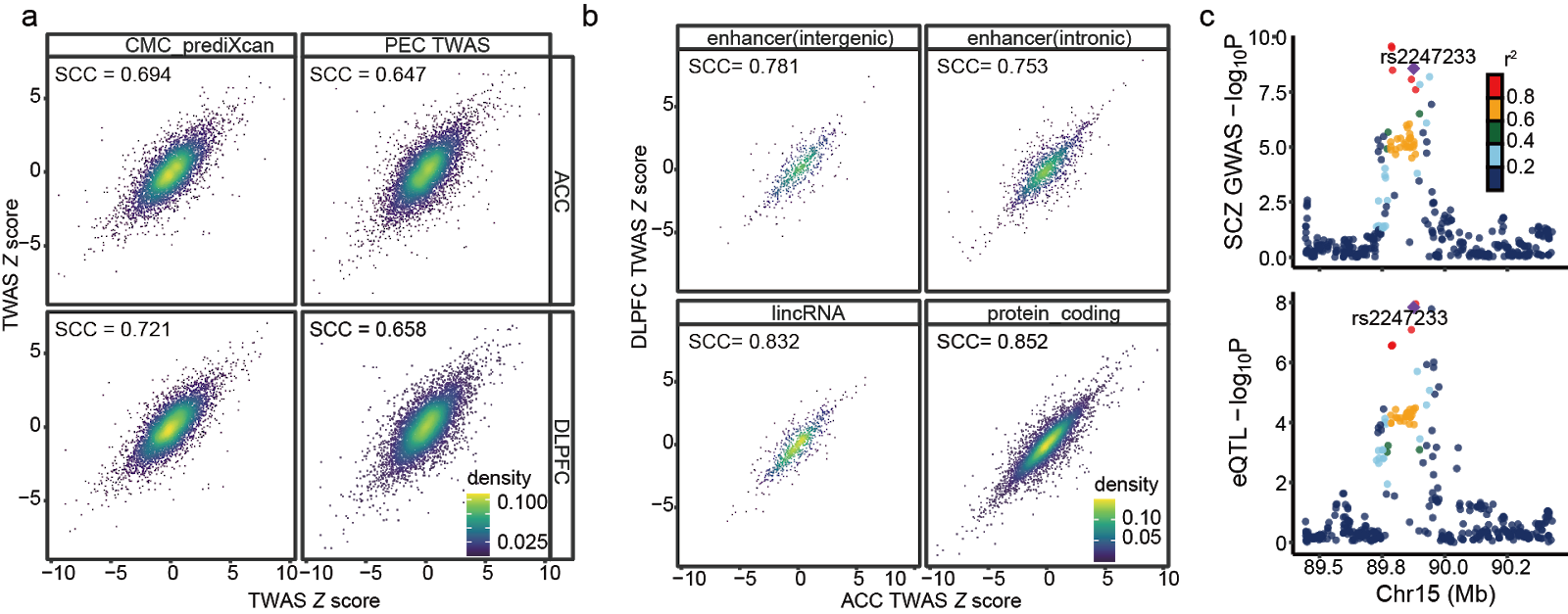

###### **Extended Data Fig. 5 | SCZ TWAS. a**, Gene TWAS Z scores compared to published reports[^32,33^](https://sciwheel.com/work/citation?ids=6164734,7359961&pre=&pre=&suf=&suf=&sa=0,0). SCC represents the Spearman correlation coefficient (ρ)(all p<10^-16^). **b**, For different types of transcripts, the TWAS Z scores between the two brain regions. SCC represents the Spearman correlation coefficient (ρ)(all p<10^-16^). **c**, Aligned Manhattan plots of SCZ GWAS and EeQTLs at the enh41216 locus generated by LocusCompare. SNPs are colored by LD (r^2^) with the lead EeQTL (rs3247233).

### Supplementary Tables

Table S1 identified the TEn list

Table S2 selected covariates for each assay

Table S3 Differential analysis results between neuronal and non-neuronal cells of different assays

Table S4 gene-enhancer link summary

Table S5 Statistics for Significant eQTLs, top SNP per transcript

Table S6 TWAS fine-mapped transcripts and the enhancer-linked genes
